# Ceramide-rich extracellular vesicles as pathogenic biomarkers in traumatic brain injury

**DOI:** 10.64898/2026.04.01.715607

**Authors:** Zainuddin Quadri, Zhihui Zhu, Xiaojia Ren, Simone M. Crivelli, Liping Zhang, P.D. Kunjadia, Patrick G. Sullivan, Bibi B. Broome, Tritia R. Yamasaki, Erhard Bieberich

## Abstract

Extracellular vesicles (EVs) contribute to the damage caused by traumatic brain injury (TBI) and can cross the blood-brain barrier (BBB). We analyzed plasma-derived EVs from human TBI patients to identify factors potentially contributing to TBI pathology. EVs were isolated using membrane affinity (ExoEasy) and size exclusion chromatography (iZone), both yielding CD9(+) and CD63(+) EVs with minimal contamination by serum albumin and apolipoprotein. Immunoblotting detected GFAP in TBI but not control EVs, indicating astrocyte-derived EVs crossing the BBB. Proteomic analysis and immunoblotting of EVs from TBI samples identified C-reactive protein and 14-3-3 proteins, which were not detected in control EVs, indicating inflammation associated with TBI. Lipidomic analysis showed ceramide enrichment in TBI EVs, validated by anti-ceramide immunoprecipitation. In a mouse closed head-controlled cortical impact model, brain EVs similarly showed elevated ceramide, confirming ceramide-rich EV release after TBI. Immunocytochemistry localized acid sphingomyelinase (ASM), a ceramide-generating enzyme, to ependymal cilia, suggesting these sites as a potential source of EVs. This was further supported by the detection of ASM in both brain- and plasma-derived EVs, along with the ciliary marker Arl13b in the brain. To assess function, we treated murine neuronal (N2a) cells with TBI EVs. Transcriptomics and STRING analyses revealed enrichment of mitochondrial-associated transcripts. Immunoblotting showed increased p53 and voltage-dependent anion channel 1 (VDAC1), which mediate ceramide-induced apoptosis. Seahorse assays showed that TBI EVs suppressed glycolysis, as indicated by reduced ECAR, while mitochondrial respiration (OCR) remained unchanged. LDH assays further indicated that TBI EVs were more neurotoxic than control EVs. Together, these findings identify ceramide-rich EVs as plasma biomarkers of TBI-induced inflammation, potential mediators of neuronal mitochondrial dysfunction, and pharmacological targets to prevent TBI-induced damage.

## Introduction

Traumatic brain injury (TBI) is a major cause of death and long-term disability, leading to both acute neuronal damage and chronic neurodegeneration. Its complex pathology involves primary mechanical injury and secondary processes that disrupt neuronal, glial, and vascular function. Beyond the primary mechanical insult, the secondary injury processes, including neuroinflammation,[27] mitochondrial dysfunction,[71, 74] and blood–brain barrier (BBB) breakdown,[6] evolve over hours to weeks post-injury and significantly influence patient outcomes.[44] Although clinical tools such as the Glasgow Coma Scale (GCS) provide valuable information on injury severity, they have limited predictive accuracy for long-term outcomes.[36, 67, 69] This underscores the urgent need for robust, mechanism-based biomarkers that can improve diagnosis, stratify severity, and guide therapeutic interventions.

Extracellular vesicles (EVs) are lipid bilayer–enclosed nanoparticles secreted by virtually all cell types, lacking a functional nucleus and replicative capacity. They serve as mediators of intercellular communication by transporting proteins, lipids, metabolites, and nucleic acids in a protected, stable form. EVs can cross the BBB, allowing brain-derived EVs to be detected in peripheral biofluids such as blood, offering a minimally invasive window into central nervous system (CNS) pathology and EV-associated pathogenic factors.[48, 60, 61]

Neuroinflammation and neuronal injury are defining features of TBI,[15, 44, 63, 78] and both have been linked to alterations in ceramide levels,[5, 19, 22] a bioactive lipid that plays a critical role in EV biogenesis.[60, 70] Ceramides involved in apoptosis,[56, 59] membrane dynamics,[3] and mitochondrial dysfunction[20, 24, 61, 66] are enriched in EVs from patients with Parkinson’s[41] and Alzheimer’s disease.[18] Elevated ceramide levels are closely linked to the activity of acid sphingomyelinase (ASM), a lysosomal enzyme that hydrolyzes sphingomyelin to ceramide.[11, 30, 38, 39, 61] We have shown that increased ASM activity induces the production of ceramide-rich EVs that target mitochondria in neurons.[61] While the effect of TBI-induced ceramide-rich EVs in neurons is not known, other studies have shown that TBI-induced EVs can reprogram gene expression in recipient cells[33] and increase the abundance neurotoxic complement proteins of the plasma-derived EVs.[29] Thus, we hypothesized that TBI induces ceramide-rich EVs that act as biomarkers of injury and actively propagate damage by reprogramming target cells at both molecular and functional levels.

In this study, we combined proteomic, lipidomic, and functional analyses of plasma-derived EVs from human TBI patients and a mouse model of controlled cortical impact. We identified glial and inflammatory markers such as GFAP and C-reactive protein (CRP), as well as 14-3-3 protein unique to TBI EVs. In addition, we found that TBI-induced EVs are enriched with ceramide and alter transcripts in recipient neuronal cells, including transcripts associated with the function of mitochondria, consistent with impaired glycolysis and increased neurotoxicity of TBI EVs. These findings establish ceramide-rich EVs as candidate biomarkers and mediators linking acute injury to altered mitochondrial function and chronic neurodegeneration, underscoring their potential for EV-based diagnostics and therapeutics.

## Materials and methods

### Animals and experimental groups

C57BL/6 male mice (8–12 months; n = 6 per group) were maintained under a protocol approved by the University of Kentucky Institutional Animal Care and Use Committee (IACUC; AUP #2017-2677). For the closed head injury (CHI) model, mice were anesthetized with isoflurane (4% induction, 3% maintenance) and positioned on a stereotaxic frame with zygomatic stabilization. Ophthalmic ointment was applied, and the scalp was disinfected and incised to expose the skull. TBI was induced using a pneumatic cortical impactor (TBI-0310; Precision Systems and Instrumentation) with a 5 mm tip, delivering 2.0 mm depth at 3.5 m/sec for 500 msec, aligned to the lambda suture. Incisions were closed with staples, and animals recovered on a heating pad. Sham animals underwent identical procedures without impact. Animals were sacrificed 48 h post-injury as described previously.[34]

### Tissue and serum collection

Right brain hemispheres were processed for EV isolation; left hemispheres were fixed in 4% PFA for immunohistochemistry. Prefrontal cortex samples were collected for biochemical analyses. Blood was obtained from the posterior vena cava, allowed to clot for 1 h at room temperature, centrifuged at 1,000 × g for 10 min, and the supernatant re-centrifuged at 3,000 × g for 15 min. Serum was stored at −80°C until use.

### Isolation of EVs from human blood plasma, mice blood serum, and mouse brain

Human blood plasma samples from both TBI patients and non-TBI controls were generously provided by the Neurobank as part of the Neuroscience Research Priority Area at the University of Kentucky, Lexington, KY. All participants were male, with an average age ranging from 28 to 30 years in both groups. Blood sample was collected from TBI and non-TBI (control) patients within 24 to 48 h post-concussion. All but one of the TBI patients were diagnosed with severe TBI, with Glasgow Coma Scale scores ranging from 3 to 5 (refer to Supplementary Table 1 for GCS and patients’ information) as noted at the time of arrival at the UK Medical Center. Blood samples were collected in EDTA-containing tubes as an anticoagulant and centrifuged at 1,500 × g for 10 min at RT to separate the plasma and aliquots then stored at −80°C. All participants provided written informed consent prior to enrollment in this IRB approved study protocol, in accordance with institutional and ethical guidelines. Demographic characteristics, including age, sex, and race, for both TBI patients and healthy control subjects are summarized in Table 1.

**Table 1.** Demographic and clinical characteristics of patients, including sex, TBI severity, age, and race.

| Subject Number | Head Trauma | Sex | Age | GCS | Blood | Race |
| --- | --- | --- | --- | --- | --- | --- |
| NSB0785 | TBI | M | 30 | 5 | venous blood/48 hours | white, non-Hispanic |
| NSB0783 | TBI | M | 23 | 4 | venous blood/48 hours | white, non-Hispanic |
| NSB0737 | TBI | M | 30 | 3 | venous blood/24 hours | white, non-Hispanic |
| NSB0661 | TBI | M | 33 | 11 | venous blood/24 hours | black or African-American, non-Hispanic |
| NSB0559 | TBI | M | 24 | 4 | venous blood/48 hours | white, non-Hispanic |
| NSB0628 | TBI | M | 42 | 3 | arterial/48 hours | white, non-Hispanic |
| NSB0036 | Control | M | 34 | N/A | Venous | white, non-Hispanic |
| NSB0747 | Control | M | 20 | N/A | Venous | native American, non-Hispanic |
| NSB0316 | Control | M | 50 | N/A | Venous | White not Hispanic |
| NSB0808 | Control | M | 18 | N/A | Venous | White not Hispanic |
| NSB0178 | Control | M | 21 | N/A | Venous | White not Hispanic |

EVs from mouse brains and blood plasma or serum were isolated as previously described.[18, 24] Briefly, mouse brains were weighed and transferred to 15 mL conical tubes. To each tube, 2 mL of Hibernate A (GIBCO, Cat# A12475-01) supplemented with 100 µL of Papain (Sigma, Cat# P3125) was added. Tubes were incubated in a 37°C water bath for 20 min with occasional shaking and intermittent pipetting to dissociate tissue and facilitate EV release by disrupting cell-to-cell connections. After 20 min, 20 µL of Protease Inhibitor Halt Cocktail (Thermo Fisher, Cat# 78442) and 6 mL of ice-cold Hibernate A were added. Tissue was further dissociated by gentle pipetting and the suspension was centrifuged at 300 × g for 5 min at 4°C to remove cells. The resulting supernatant was centrifuged at 4,000 × g for 20 min to eliminate larger debris. This was followed by ultracentrifugation using an SW 41 Ti rotor at 10,000 × g for 40 min at 4°C. The final supernatant was processed using the ExoEasy Maxi Kit (Qiagen, Cat# 76064) according to the manufacturer’s instructions to isolate EVs.

Similarly, EVs from mouse blood serum or human blood plasma were isolated using the ExoEasy Maxi Kit (Qiagen, Cat# 76064) following a similar protocol. To assess the purity of EVs, size exclusion chromatography was performed using the qEVsingle iZon Gen2 column following the manufacturer’s instructions. Briefly, the column was equilibrated by flushing twice with 3 mL of PBS. Human blood plasma was pre-cleared by centrifugation at 1,500 × g for 10 min, followed by an additional centrifugation at 10,000 × g for 10 min. The resulting supernatant was loaded onto the column. A total of seven fractions (~200 µL each) were collected, and EVs were quantified using nanoparticle tracking analysis (NTA) with the ZetaView system. Based on particle concentration, fractions 4 through 7 were selected for immunoblot analysis.

### LDH neurotoxicity assay

EVs were added to Neuro-2a (N2a) cells (ATCC (CCL-131™) to quantify the cytotoxicity of EVs as described previously.[61] Briefly, one day prior to incubation after reaching 70-80% cell confluence, the cells were washed with PBS, and cells were incubated with serum-free DMEM. Next day, 10,000 EVs per cells were added and 24 h post incubation, 50 μl media were collected and LDH measurement was performed as per manufacturer’s protocol using CyQUANT™ LDH Cytotoxicity Assay Kit (Invitrogen, Cat# C20301). Optical density measurement was performed by measuring the absorbance at 490 and 680. The % neurotoxicity was calculated using the formula % Cytotoxicity = (Compound-treated LDH activity - Spontaneous LDH activity) / (Maximum LDH activity - Spontaneous LDH activity) x100), and the graph was plotted as fold change.

### Transcriptomics analysis

As outlined in the previous methods section, an equal number of EVs isolated from the brains of either sham or TBI mice were added to serum-free N2a cells. After 24 h of incubation, the cells were washed with PBS and then harvested. The cells were homogenized using a QIAshredder (Qiagen, Cat#79654), and RNA was isolated using RNeasy Plus Mini Kit (Qiagen, Ref#74134). RNA samples were submitted to Novogene (https://en.novogene.com) for quality assessment and RNA sequencing analysis. Novogene conducted RNA sequencing and performed initial statistical analysis using DESeq2 to identify differentially expressed genes.

### Proteomics analysis

An equal number of human blood plasma-derived EVs from TBI and control groups was denatured and loaded on 4–20% SDS-PAGE gels and stained overnight with Coomassie Brilliant Blue R-250 (BIO-RAD, Cat#161-0436). Gels were subsequently destained in a solution containing 10% acetic acid, 50% methanol, and 40% H O until the background was clear. Protein bands at approximately 25 kDa from both TBI- and control-derived EVs were excised using a sterile blade and submitted to Proteomics and Mass Spectrometry Core Laboratory, Augusta University, Augusta GA for shotgun proteomics analysis. LC-MS/MS was performed on an Orbitrap Fusion Tribrid mass spectrometer (Thermo Scientific, New York, NY, USA) coupled to an Ultimate 3000 nano-UPLC system (Thermo Scientific), following protocols previously described by our group.[25]

### Immunoblot

An equal number of EVs isolated from control (non-TBI) and TBI human plasma samples was mixed with 5× SDS sample buffer (50 mM Tris-HCl, pH 6.8; 4% SDS; 25% glycerol; 0.1% bromophenol blue; 5% 2-mercaptoethanol) and denatured at 95°C for 10 min. The denatured proteins were resolved by SDS-PAGE using a 4-20% Precast Mini-PROTEAN TGX gel (Bio-Rad, Cat# 4561094) and transferred onto a nitrocellulose membrane via electrophoresis for 1 h. Following transfer, membranes were stained with Ponceau S (Sigma, Cat# P7170) to verify protein transfer, then blocked with 5% non-fat dry milk for 1 h to prevent non-specific binding. Membranes were incubated overnight at 4°C with the appropriate primary antibodies. After three washes with 0.07% Tween 20 in Tris-buffered saline (TBS-T), membranes were incubated with HRP-conjugated secondary antibodies for 2 h at RT. Immunoreactive bands were visualized using either SuperSignal West Pico or West Femto chemiluminescent substrates and imaged using the Azure 300 Chemiluminescent Imaging System. The primary antibodies used in this study were anti-CD9 (CBL162, Millipore Sigma), Anti-CD63 (Invitrogen, Cat#MA530187), anti-CD81 (BioScience, Cat#EV-hCD81-A-555), anti-Flotillin 2 (BD Transduction laboratories, Cat#610383), anti-Annexin A1 (Abcam, Cat# ab214486), anti-Acid Sphingomyelinase (Proteintech, Cat#14609-1-AP), anti-Cox4I2 (Proteintech, Cat#14463-1-AP), anti-GAPDH (Proteintech, Cat#HRP-60004), anti-p53 (Cell signaling, Cat#3253T), anti-β-Actin (Santa Cruz, Cat# sc-47778), anti-VDAC1 (Santa Cruz, Cat#sc-390006), anti-Human Serum Albumin (R&D System Cat#MAB1455), anti-C-Reactive Protein (Proteintech, Cat#24175-1-AP), anti-GFAP (Dako, Cat#Z0334), anti-GFAP (Novus Biologicals NB300-141).

### Targeted lipidomics (LC-MS/MS analysis)

The lipidomic analysis was performed on the EVs and the mouse brain lysate. EVs for lipidomics were isolated from mice brain, mice blood serum and human blood plasma, whereas the brain lysate was obtained from a piece of prefrontal cortex homogenized in homogenization buffer containing 50 mM Tris-HCl, 0.5 mM EDTA, 0.5 M Sucrose, and 25mM KCl. The amount of ceramide content of EVs or brain lysate was measured by the lipidomics core facility at the Medical University of South Carolina, SC (https://hollingscancercenter.musc.edu/) as described previously.[8, 9, 16] The values were normalized to total phosphate (Pi) in the lipid fraction, total EV count, or total protein content (µg) as indicated in the legends to the pertinent figures.

### Immunoprecipitation of EVs from human blood plasma using an anti-ceramide antibody

Immunoprecipitation of EVs using anti-ceramide rabbit IgG was performed as previously described.[24] Protein A/G Sepharose magnetic beads (Pierce, Cat# 88803) were washed three times with PBS using a magnetic separator and non-specific binding sites blocked with 1% bovine serum albumin (BSA; Sigma, Cat# A2153) for 1 h at RT. Beads were then incubated with equal amounts of either anti-ceramide antibody or control IgG in 0.5% BSA/PBS and rotated at RT for 2 h to facilitate antibody binding. Separately, equal numbers of EVs isolated from TBI and control samples were pre-incubated with 20 µL of human Fc receptor (FcR) blocking reagent (Miltenyi Biotec, Cat# 130-059-901) on ice for 30 min to minimize non-specific binding. The pre-blocked EVs were then incubated with beads and the mixture was rotated at 4°C for 2 h. Beads were separated using a magnetic separator, and the unbound fraction was collected and subsequently incubated with ceramide or IgG-conjugated beads at 4°C under continuous rotation for immunocapture. The next day, beads were separated magnetically, and the supernatant was collected as flow-through. Beads were then washed ten times with PBS (10 min each wash on a shaker) to ensure thorough removal of non-specifically bound material. To lyse the bead-bound EVs, 100 µL of RIPA buffer (50 mM Tris-HCl pH 7.4, 150 mM NaCl, 1% NP-40, 0.1% SDS, 1 mM EDTA, supplemented with protease inhibitors) was added, and samples were incubated on ice for 10 min. Lysates were centrifuged at 14,000 × g for 1 min at 4°C to separate proteins from the beads. Supernatants were collected, mixed with 5× SDS loading buffer containing β-mercaptoethanol (50 mM Tris pH 6.8, 4% SDS, 25% glycerol, 0.1% bromophenol blue), and 30 µL of each lysate was loaded onto an SDS-PAGE gel for CRP and CD9 detection via immunoblotting.

### EV immobilization on coverslips and immunolabeling

EVs were immobilized on coverslips following a previously described protocol.[12] To render the surface of glass coverslips (FisherBrand, Cat# 1254580) hydrophilic, they were submerged in a freshly prepared solution of deionized (DI) water, hydrogen peroxide (H□O□, Cat#216763), and ammonium hydroxide (NH OH; Thermo Scientific, Cat# 42330-5000) mixed in a 5:1:1 ratio. The solution was heated to 88°C and coverslips were incubated in this solution for 10 min. After treatment, coverslips were thoroughly rinsed with DI water 3–5 times. The cleaned coverslips were transferred to a 24-well plate and incubated with 5% (v/v) APTES (3-aminopropyltriethoxysilane; Thermo Scientific, Cat# 42330-5000) in 95% ethanol for 10 min at RT to promote silanization and introduce primary amine groups onto the silica surface, facilitating EV immobilization. Following incubation, coverslips were rinsed three times with DI water and further incubated with 1% (v/v) glutaraldehyde (Sigma-Aldrich, Cat# G7651) in DI water for 1 h at RT. Coverslips were then rinsed three additional times with DI water. For EV immobilization, samples were diluted in PBS to a final concentration of 10 particles per 50 µL. A 50 µL aliquot of the EV suspension was added to each functionalized coverslip in a humidified chamber and incubated for 1 h at RT. Unbound EVs were removed by gentle washing with PBS three times for 10 min each on a shaker. Subsequently, immobilized EVs were fixed using a mixture of 2% paraformaldehyde (PFA) and 0.1% glutaraldehyde in PBS for 20 min at RT. Coverslips were again washed three times with PBS for 10 min each. To block nonspecific binding, coverslips were incubated with Tris-ethanolamine buffer (0.1 M Tris, 50 mM ethanolamine; Sigma-Aldrich, Cat# E9508), pH 9.0, for 30 min with gentle shaking, followed by three washes with PBS for 10 min each. Coverslips were then incubated with 0.05% casein (Sigma, Cat#C7078) in PBS for 90 min with shaking. After blocking, coverslips were ready for immunolabeling with anti-CD9 and anti-CRP antibodies following the immunocytochemistry protocol.

### Seahorse analysis

N2a cells were seeded in a Seahorse XFe96 Standard Cell Culture Microplate (Agilent, Cat# 102416-100) and cultured until they reached 70–80% confluency. One day prior to treatment, the cells were incubated with serum-free medium. The following day, EVs were added at a concentration of 5,000 EVs per cell. After 24 h of incubation with EVs, the metabolic and bioenergetic profiles of N2a cells were assessed using the Seahorse XF Cell Mito Stress Test on a Seahorse XFe96 Analyzer (Agilent Technologies, North Billerica, MA). Before the assay, the culture media were replaced with Seahorse XF assay medium, and a standard mitochondrial bioenergetics and glycolytic metabolic test was performed as described previously.[17, 35] The resulting data were normalized to cell numbers using the BioTek Cytation 5 imaging system (Agilent).

### Statistical analysis

GraphPad Prism version 10.2 was used for statistical analysis. An unpaired t-test was used to compare differences between two groups, and a one-way ANOVA was applied for comparisons among three groups. Statistical significance was set as p <0.05 and star significance were distributed as * for *p* < 0.05, ** for *p* < 0.01, *** for *p* < 0.001, and **** for *p* < 0.0001.

## Results

### TBI-induced EVs from human plasma contain protein markers of neuronal injury and inflammation

To isolate EVs from human blood plasma, two different methods were employed: an affinity-based approach using the ExoEasy Maxi Kit (Qiagen), and a size exclusion chromatography (SEC) method utilizing qEV columns (iZon). Nanoparticle tracking analysis (NTA; Zetaview) revealed that the concentration of nanoparticles isolated from TBI plasma samples using ExoEasy was significantly higher (1.5-fold) than those from control samples (Fig. 1A). Particle size distribution analysis showed that the majority of nanoparticles were EVs of approximately 150 nm in diameter (Fig. 1A). Western blot characterization confirmed the presence of canonical EV markers, including Flotillin-2, tetraspanins (CD9, CD63, CD81), and Annexin A1 (Fig. 1B). To assess the purity of the EV preparations, we also probed for potential protein contaminants such as Human Serum Albumin (HSA), Apolipoprotein A1 (ApoA1), and Calreticulin. These proteins were undetectable in EVs isolated via the ExoEasy method, indicating a high level of purity (Fig. 1C). To further validate the purity and confirm the EV-containing fractions, we performed SEC-based EV isolation using iZon columns, collecting seven consecutive 200 µL fractions. NTA data indicated that fractions 5–7 contained the majority of particles, which were subjected to immunoblotting for EV markers CD9 and CD63 (Supplementary Fig. 1A). ApoA1 and HSA were not detected in any of the EV fractions, verifying EV purity. The particle size of EVs in fractions 5-7 ranged from 100 to 150 nm, comparable to the size of EVs isolated with ExoEasy. However, ExoEasy-based isolation produced approximately 5-fold greater EV yields than SEC-based isolation at comparable purity, establishing this as the preferred method for EV-intensive downstream analyses, including proteomics, lipidomics, and functional assays.

**Figure 1.**
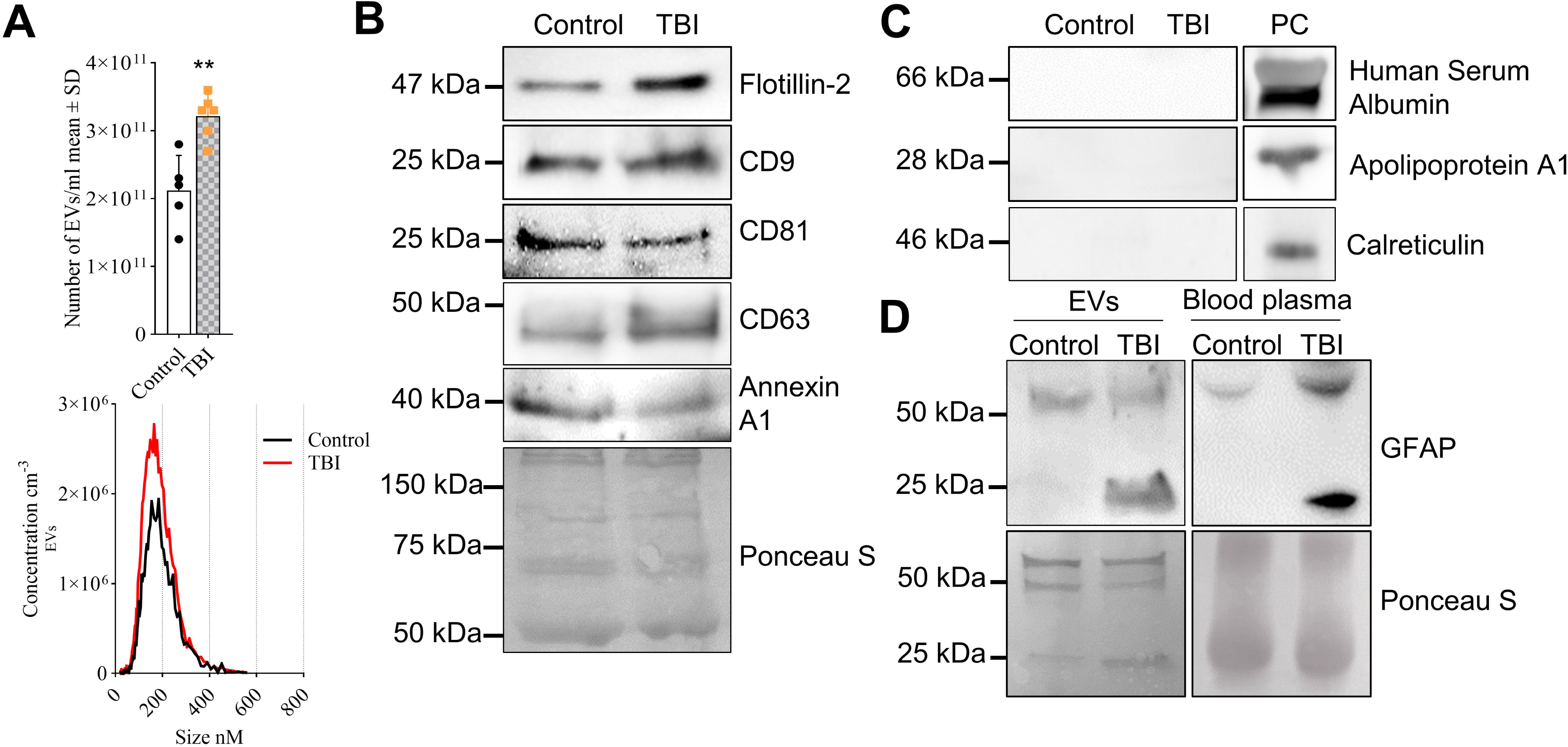
Plasma-derived EVs from TBI patients are enriched with GFAP. (**A**) NTA of EVs: concentration (top) and particle size distribution curve (bottom). Mean ± SD, unpaired t-test, *n* = 5 for control and n=6 for TBI, * *p* < 0.05, ** *p* < 0.01, *** *p* < 0.001, and **** for *p* < 0.0001. (**B**) Immunoblot of canonical EV-associated proteins Flotillin-2, CD9, CD81, CD63 and Annexin A1. Blot was stained with Ponceau S for loading control. (**C**) Immunoblot of a non–EV-associated protein Human Serum Albumin, Apolipoprotein A1 and Calreticulin, which are detected only in plasma control (PC). (**D**) Immunoblot of GFAP in plasma (right panel) and plasma-derived EVs (left panel) (top) with Ponceau S staining (bottom) as loading control.

To determine the origin of TBI-induced plasma EVs, we performed immunoblotting for Glial Fibrillary Acidic Protein (GFAP), a brain-specific cytoskeletal protein and a biomarker for intracranial injury, known to be elevated in the blood following TBI.[1, 10, 42] Moreover, truncated breakdown products of GFAP have been reported in plasma post-TBI.[57] In addition to the ~50 kDa full-length protein, a truncated GFAP band (~25 kDa), consistent with proteolytic cleavage products previously reported in TBI, was exclusively detected in TBI EVs and blood plasma (Fig. 1D), indicating that astrocytes secreted truncated GFAP associated with EVs in response to brain injury.

Interestingly, Ponceau S staining of EV protein blots consistently showed a distinct band at ~25 kDa in TBI samples, which was less intense and appears slightly higher in molecular weight in controls (Fig. 1D). To identify the protein, EV samples were subjected to Coomassie staining (Fig. 2A), and the ~25 kDa band was excised for mass spectrometry. Proteomic analysis revealed several candidates (see Supplementary Table 1), among which two proteins, 14-3-3 and C-reactive protein (CRP), were highly enriched in TBI EVs (Fig. 2B and C). 14-3-3 protein is highly abundant in the brain,[49] whereas its detectable levels in serum or plasma are generally low and depend on blood processing conditions.[32] C-reactive protein (CRP) is a well-established marker of systemic inflammation and previously reported in TBI plasma.[68, 72, 76] Immunoblotting confirmed the presence of 14-3-3 protein and CRP exclusively in TBI-derived EVs (Fig. 2B and C).

**Figure 2.**
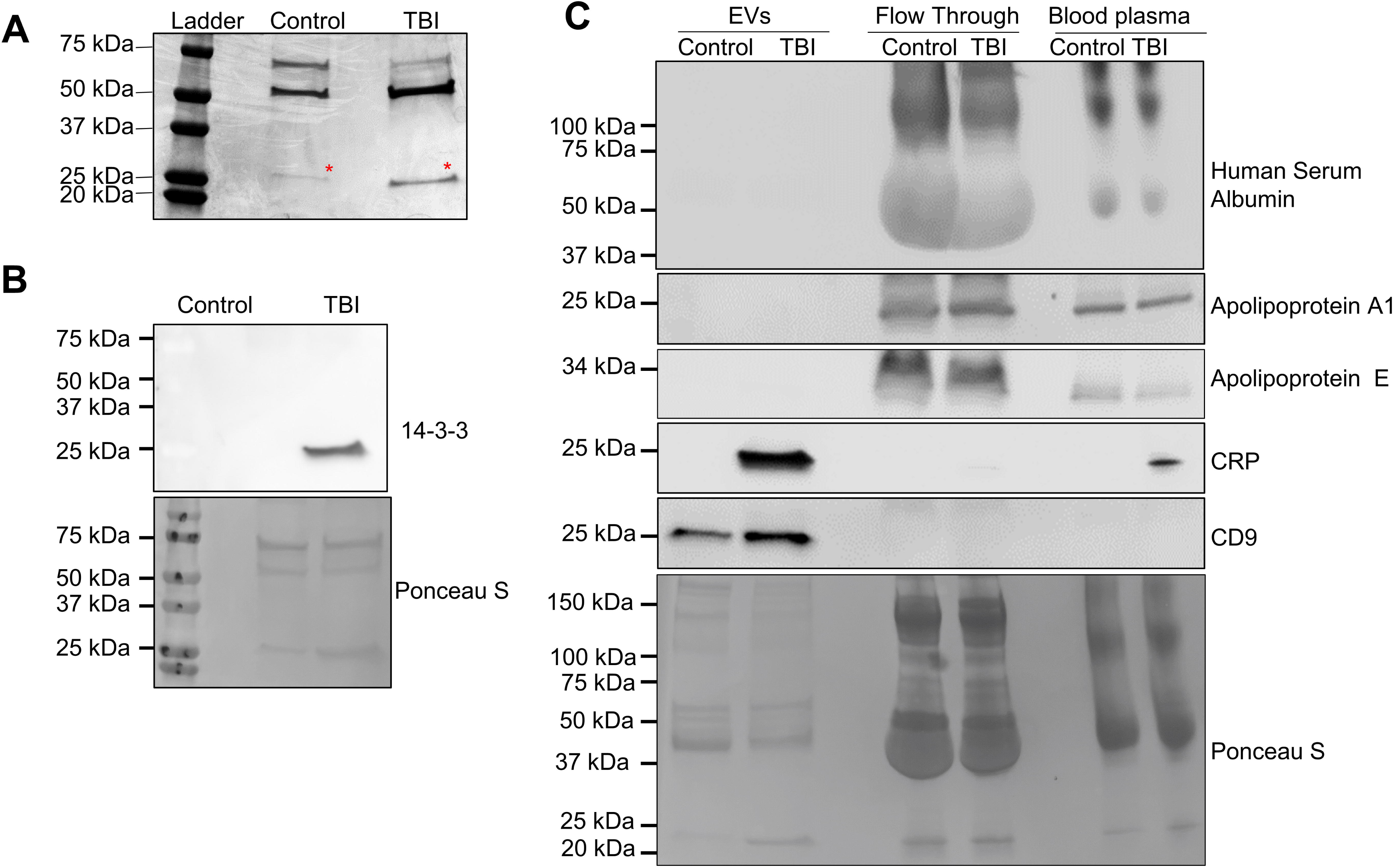
TBI EVs contain 14-3-3 protein and CRP. (**A**) EV protein from human control and TBI plasma was separated by SDS-PAGE. The red asterisk denotes a ~25 kDa Coomassie-stained band, which was excised and subjected to proteomic analysis. (**B**) Immunoblot of 14-3-3 in plasma-derived EVs from TBI and control samples (top). Ponceau S staining shown as loading control (bottom). (**C**) Immunoblot for CRP, EV (CD9)- and non–EV (Human Serum Albumin, Apolipoprotein A1 and Apolipoprotein E)-associated proteins in EVs fraction, flow-through fraction, and whole plasma; Ponceau S staining as loading control (bottom).

To determine whether CRP was associated with EVs or represented a protein contaminant in the EV fraction, we performed comparative immunoblotting of EVs isolated using ExoEasy and the corresponding flow-through (non-vesicular fraction), both normalized to equivalent plasma input. CRP was undetectable in the flow-through, suggesting the majority of TBI-induced plasma CRP was bound to EVs (Fig. 2C). As additional controls, other potential protein contaminants such as ApoA1, ApoE, and HSA were detected in the flow-through but were absent from the EV fraction, supporting the specificity and purity of the isolated EVs. The expression levels of the low molecular weight GFAP band and CRP were positively correlated with TBI severity and showed an inverse relationship with the Glasgow Coma Scale score, indicating the association of EV-bound GFP and CRP with TBI severity (Supplementary Fig. 2).

To test if CRP colocalized with CD9(+) EVs, human blood plasma EVs were immobilized on coverslips and immunolabeled with anti-CD9 and with anti-CRP antibody. Supplementary Figure 3 shows that the proportion of EVs co-labelling for CD9 and CRP was 31.4-fold higher in TBI-EVs than controls, consistent with the results from immunoblotting of isolated EVs. Taken together, the association of 14-3-3 protein, GFAP, and CRP with EVs in plasma indicates that TBI induces secretion of EVs as part of an inflammatory process, at least in part originating in astrocytes and detectable as EV-associated plasma biomarkers.

### Ceramide enrichment is characteristic for TBI-induced EVs in human plasma and mouse serum and brain

Our previous studies showed that neuroinflammation increases ceramide in astrocytes,[80] which secrete ceramide-rich EVs in Alzheimer’s disease (AD). Others have reported that ceramide generation is induced by TBI as well.[5] To investigate whether EVs isolated from the human blood plasma of TBI patients carry distinct ceramide signatures, we performed targeted liquid chromatography-tandem mass spectrometry (LC-MS/MS). This lipidomic analysis revealed a significant increase in C18:1 and C20:0 ceramide species in TBI EVs compared to controls (Fig. 3A). Levels of C24:0 ceramide were reduced in TBI EVs which aligns with the previous finding of significantly increased C18:0 and decreased C24 ceramide levels in the blood plasma of post TBI mice, although EVs were not investigated in this study.[50] Additionally, the level of sphingosine-1-phosphate (Sph-1P) was significantly lower in TBI EVs relative to controls (Fig. 3A), indicating a shift in sphingolipid metabolism induced by TBI.

**Figure 3.**
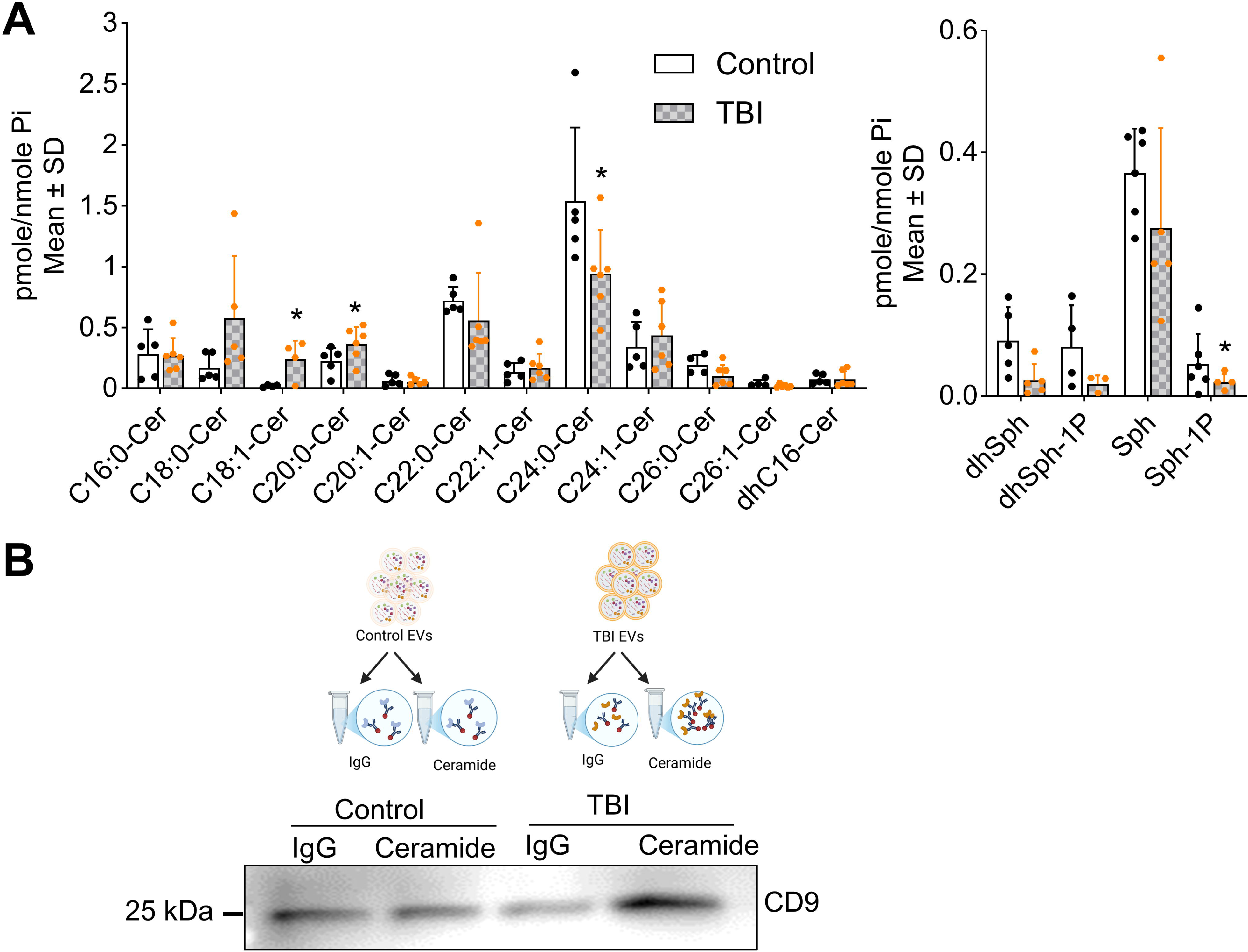
Ceramide is enriched in human TBI EVs. (**A**) Targeted sphingolipidomics (HPLC-MS/MS) of plasma-derived EVs normalized to lipid phosphate (Pi). Mean ± SD, unpaired t-test, *n*=5 for control and *n*=6 for TBI, * *p* < 0.05, ** *p* < 0.01, *** *p* < 0.001, and **** for *p* < 0.0001 (**B**) Upper panel is illustration of EV isolation from TBI and control plasma using anti-ceramide or IgG antibodies by immunoprecipitation. Lower panel for Western blot analysis of EVs immunoprecipitated with anti-ceramide antibody and probed for CD9.

To determine the proportion of ceramide-rich EVs within the population of TBI-induced plasma EVs, we performed immunoprecipitation (IP) using an anti-ceramide antibody (rabbit IgG) produced in our laboratory and IgG control immobilized on agarose beads,[40] followed by immunoblotting for CD9. In TBI-plasma-derived EV samples, IP with the anti-ceramide antibody pulled down significantly more CD9-positive EVs compared to IgG controls, while there was no increase of the CD9 signal in EVs isolated from non-TBI patients. This data indicated that ceramide-rich EVs were specifically elevated in TBI plasma (Fig. 3B).

To validate our findings in a preclinical TBI model, we tested if ceramide enrichment was also characteristic for EVs isolated from brain tissue and serum of closed head injury (CHI)-exposed mice. EV isolation followed a procedure previously published by our laboratory.[25] Lipidomic profiling revealed significant upregulation of total ceramide in both the brain cortex and brain-derived EVs following TBI (Fig. 4). Specifically, levels of C16:0, C18:0, C20:1, and C22:1 ceramides were elevated in brain tissue, accompanied by a decrease in dhSph-1P (Fig. 4A). Brain-derived EVs showed increased levels of C16:0, C18:0, C20:0, C22:0, dihydrosphingosine, and sphingosine (Fig. 4B). Moreover, C18:0 and C18:1 ceramide were consistently upregulated in mouse brain cortex, brain-derived EVs, serum EVs from TBI mice, and plasma EVs from human TBI patients, highlighting a conserved ceramide signature associated with TBI pathology.[50] (Supplementary Fig. 4).

**Figure 4.**
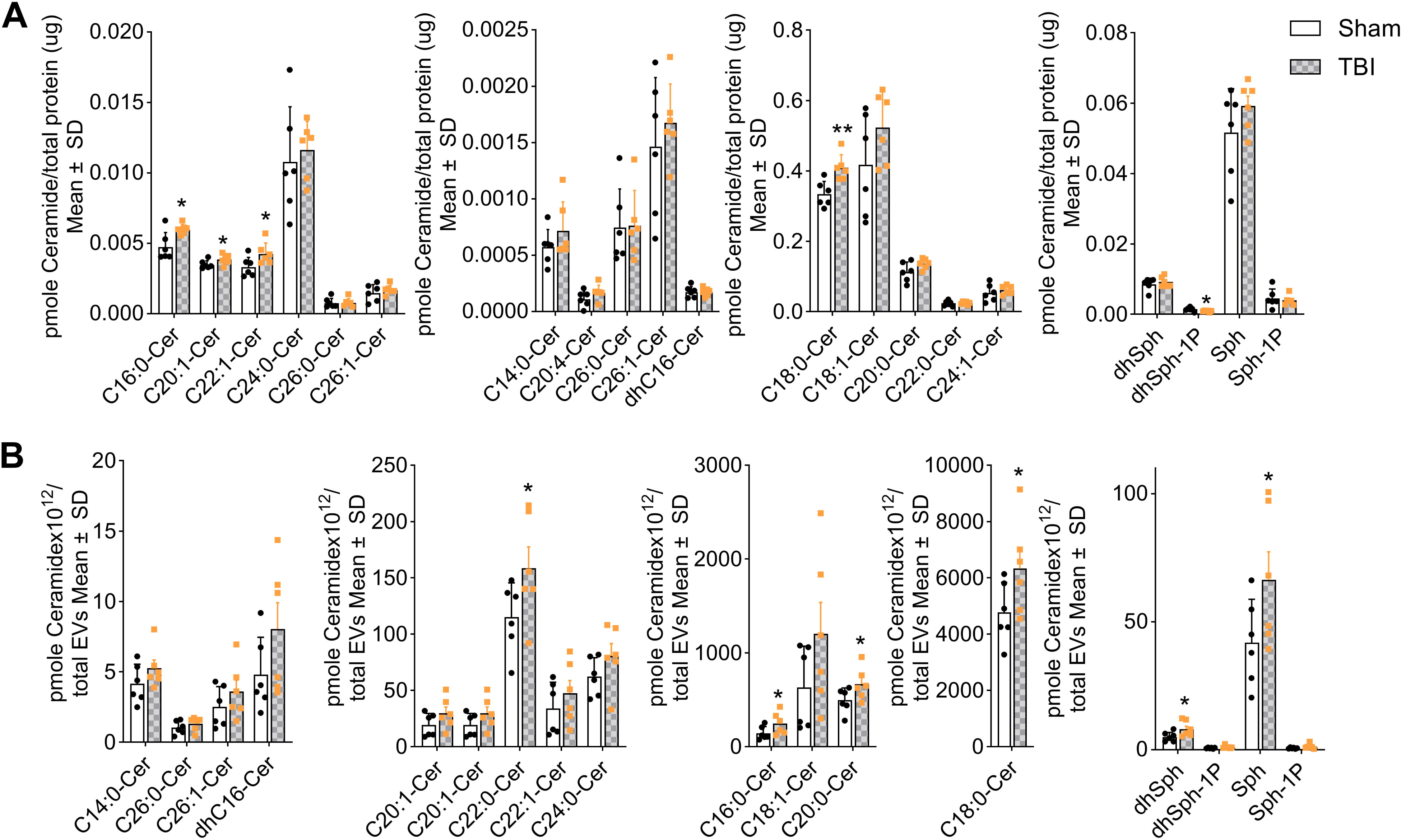
Ceramide is enriched in mouse TBI brain and EVs. Targeted sphingolipidomics (HPLC-MS/MS) of (**A**) brain lysates normalized to total protein (µg), and (**B**) brain-derived EVs normalized to total EV number. Mean ± SD, unpaired t-test, *n* = 6 for sham and TBI, * *p* < 0.05, ** *p* < 0.01, *** *p* < 0.001, and **** for *p* < 0.0001.

### TBI induces ASM translocation to ependymal cilia and increases a cilia marker in plasma EVs

Previously, we have shown that acid sphingomyelinase (ASM), a key enzyme involved in the hydrolysis of sphingomyelin to ceramide, is instrumental in the generation of ceramide-rich EVs.[18, 61] Further, ASM has been implicated in the pathophysiology of TBI.[53] To evaluate the role of ASM in our TBI model, we examined ASM expression in the brain cortex of TBI-affected mice. Immunoblotting revealed a significant increase in ASM protein levels in the brain cortex of TBI mice compared to sham (non-TBI control) (Fig. 5A), consistent with elevated ceramide production observed by lipidomics analysis (Fig. 4). While there is no data available on ASM levels in human brain after TBI, we examined whether ASM was present in plasma-derived EVs from human TBI patients. Immunoblotting showed that both, plasma EVs from non-TBI controls and TBI patients contained ASM, although there was no significant change in ASM levels in TBI EVs compared to controls (Fig. 5C). Since cilia are a source for EVs and deciliation of ependymal cells was observed in TBI,[75] we performed immunocytochemistry on brain cryosections from sham and TBI-exposed mice. We found that TBI induced translocation of ASM to ependymal cilia concurrent with punctate ceramide labeling of the ciliary membrane (Fig. 5C). Immunoblotting showed that the level of Arl13b, a ciliary marker protein, was increased in plasma EVs from human TBI patients (Fig. 5D), indicating that a proportion of ceramide-rich EVs is derived from cilia, most likely due to TBI-induced translocation of ASM to the ciliary membrane.

**Figure 5.**
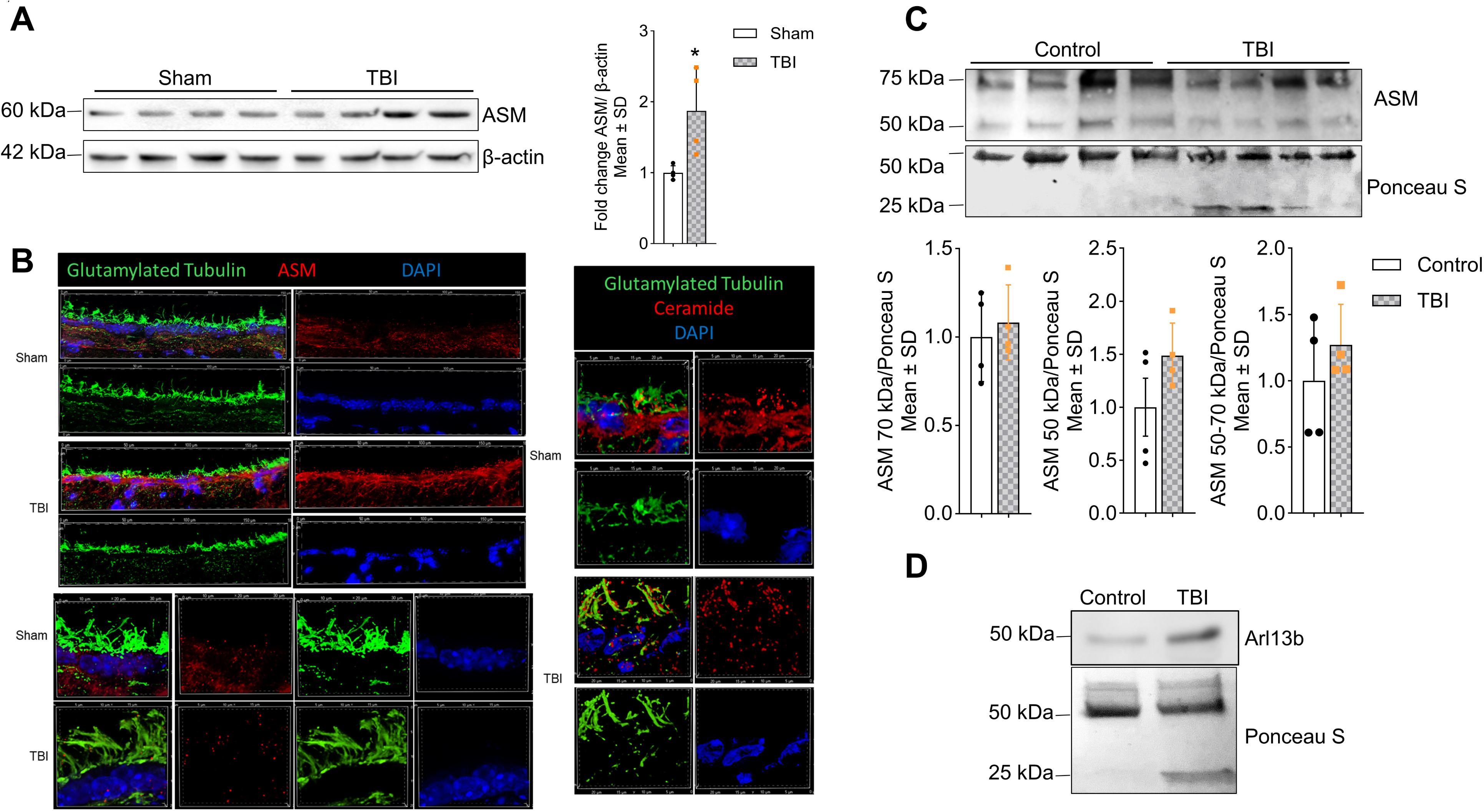
TBI EVs originate in ependymal cilia. (**A**) Immunoblot for ASM in TBI and sham mouse cortex (right) with quantification normalized to β-actin (left), Mean ± SD, unpaired t-test, *n* = 4, * *p* < 0.05, ** *p* < 0.01, *** *p* < 0.001, and **** for *p* < 0.0001. (**B**) Immunofluorescence images of mouse brain cortex showing cilia from ependymal cells of TBI and sham groups. Left panels: ependymal cells immunolabeled for ASM (red) and glutamylated tubulin, a cilia marker protein (green). Right panels: ependymal cells immunolabeled for ceramide (red) and glutamylated tubulin (green). Nuclei are counterstained with DAPI (blue). (**C**) Immunoblot for ASM in human plasma-derived EVs (top), Ponceau S staining as a loading control (middle), and quantification of ASM bands at ~70 kDa and ~50 kDa (bottom). Mean ± SD, unpaired t-test, *n* = 4, * *p* < 0.05, ** *p* < 0.01, *** *p* < 0.001, and **** for *p* < 0.0001. (**D**) Immunoblot for human plasma-derived EVs showing ciliary marker Arl13b (upper) and stained with ponceau S as loading control.

### EVs from TBI-exposed mouse brain promote enrichment of transcripts and proteins associated with mitochondrial dysfunction in N2a cells

To assess the functional impact of EVs derived from TBI-affected brain tissue on neuronal cells, we performed transcriptomic analysis after incubation of mouse neuronal (N2a) cells with EVs isolated from sham (non-TBI) vs. TBI-exposed mouse brain (see Fig. 6A for experimental outline). A control group of N2a cells that were not exposed to EVs was included and designated as ‘control’. Total RNA was isolated from each group and submitted for RNA sequencing. Transcriptomic analysis revealed distinct profiles of transcript enrichment in N2a cells exposed to TBI brain-derived EVs compared to those treated with sham EVs or left untreated as control, as shown in the heat map (Fig. 6B). Cluster analysis of significantly altered RNAs revealed enrichment in pathways related to mitochondrial oxidative phosphorylation, mitochondrial translation, and sphingolipid metabolism, highlighting a potential role of ceramide and other sphingolipids in mediating TBI-related effects (Fig. 6C).

**Figure 6.**
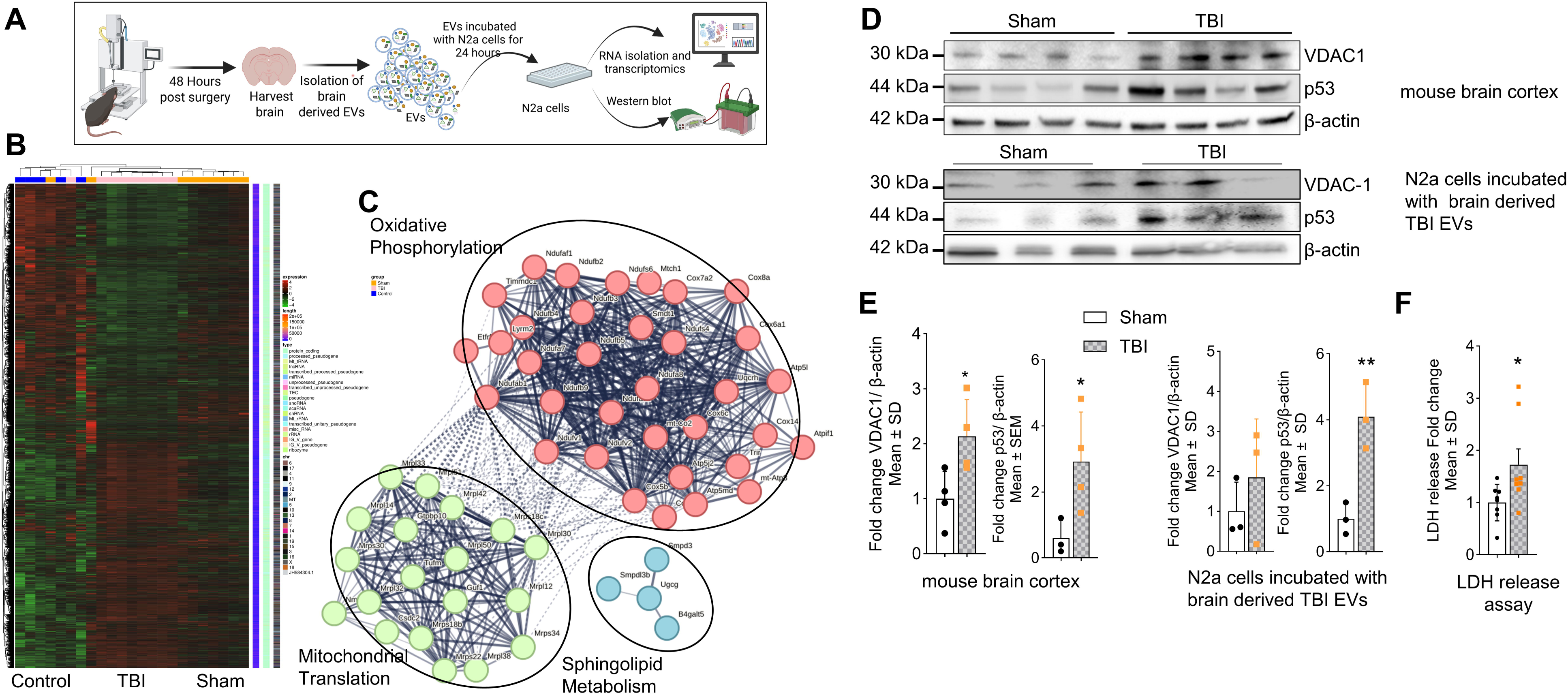
EVs from mouse TBI brain modulate expression of mitochondrial proteins in N2a cells. (**A**) Schematic illustration of TBI/sham mouse brain-derived EV isolation, incubation of N2a cells with EVs, and downstream functional analyses (transcriptomics and protein assays). (**B**) Heat map of transcriptomic changes in N2a cells incubated without EVs (control, *n* = 9), with sham EVs (*n* = 9), or with TBI EVs (*n* = 9). (**C**) STRING analysis of transcripts with *p* < 0.04 selected for cluster analysis of oxidative phosphorylation, mitochondrial translation, and sphingolipid metabolism. (**D**) Immunoblot of VDAC1 and p53 in mouse brain cortex (upper panel) and in N2a cells treated with sham or TBI EVs (lower panel) normalized to loading control β-actin. (**E**) Quantification expressed in fold-change for the blots shown above. Mean ± SD, unpaired t-test, *n* = 4 for brain cortex and *n*=3 for N2a cells, * *p* < 0.05, ** *p* < 0.01, *** *p* < 0.001, and **** for *p* < 0.0001. (**F**) LDH release assay showing fold change in N2a cell death after incubation with brain-derived EVs from TBI and sham mice. *n* = 8 for sham and *n*=9 for TBI. Mean ± SD, unpaired t-test, * *p* < 0.05, ** *p* < 0.01, *** *p* < 0.001, and **** for *p* < 0.0001.

Further functional annotation of differentially expressed genes (DEGs) demonstrated that genes involved in mitochondrial function, glycolysis, and inflammation were among the most significantly affected (Supplementary Fig. 5A–D). Notably, mitochondria-associated transcripts showed an enrichment in the TBI vs. control groups, suggesting that EVs from TBI brains disrupt mitochondrial homeostasis (Supplementary Fig. 5D). Additional DEG analysis showed that numerous transcripts were significantly altered in N2a cells exposed to TBI EVs versus sham EVs, as illustrated by fold-change distributions, the Volcano plot (Supplementary Fig. 5A–D) and log fold change dot plot (Supplementary Fig. 6), confirming a broad functional impact of TBI EVs on neuronal transcriptomes, including genes for mitochondria, glycolysis and inflammation.

We next evaluated the expression of key mitochondrial proteins to assess whether transcriptomic changes were mirrored at the protein level. In the cortex of TBI mice, expression of Voltage-Dependent Anion Channel 1 (VDAC1), a mitochondrial protein known to bind to ceramide, was significantly increased compared to sham (Fig. 6D and E).[20, 21, 24] Importantly, N2a cells incubated with TBI mouse brain-derived EVs also exhibited a moderate increase in VDAC1 expression, indicating a functional effect of EV exposure. Similarly, the tumor suppressor p53, a stress-responsive protein implicated in neuronal injury and binding to ceramide[26, 37] was elevated both in TBI brain tissue and in N2a cells treated with TBI brain-derived EVs (Fig. 6D and E). Additionally, to examine the neurotoxic potential of TBI brain-derived EVs, we measured lactate dehydrogenase (LDH) release in N2a cells following EV exposure. A significant increase in LDH release was observed in cells treated with TBI EVs (36% cell death) compared to sham EVs (23% cell death), indicating elevated cytotoxicity (Fig. 6F).

To further explore the metabolic impact of EVs isolated from human TBI plasma, we performed extracellular flux analysis using the Seahorse assay to measure oxygen consumption rate (OCR) and extracellular acidification rate (ECAR) in N2a cells (workflow in Fig. 7A). While no significant differences in OCR were observed among cells treated with (non-TBI and TBI) EVs or without EVs (control) (Fig. 7B), non-mitochondrial consumption and basal respiration was significantly reduced in the TBI EV group compared to the control group (Fig. 7C). Strikingly, glycolytic function, as measured by ECAR, was significantly decreased in N2a cells exposed to human TBI EVs compared to non-TBI EVs (Fig. 7D), indicating that TBI EVs impair glycolysis.

**Figure 7.**
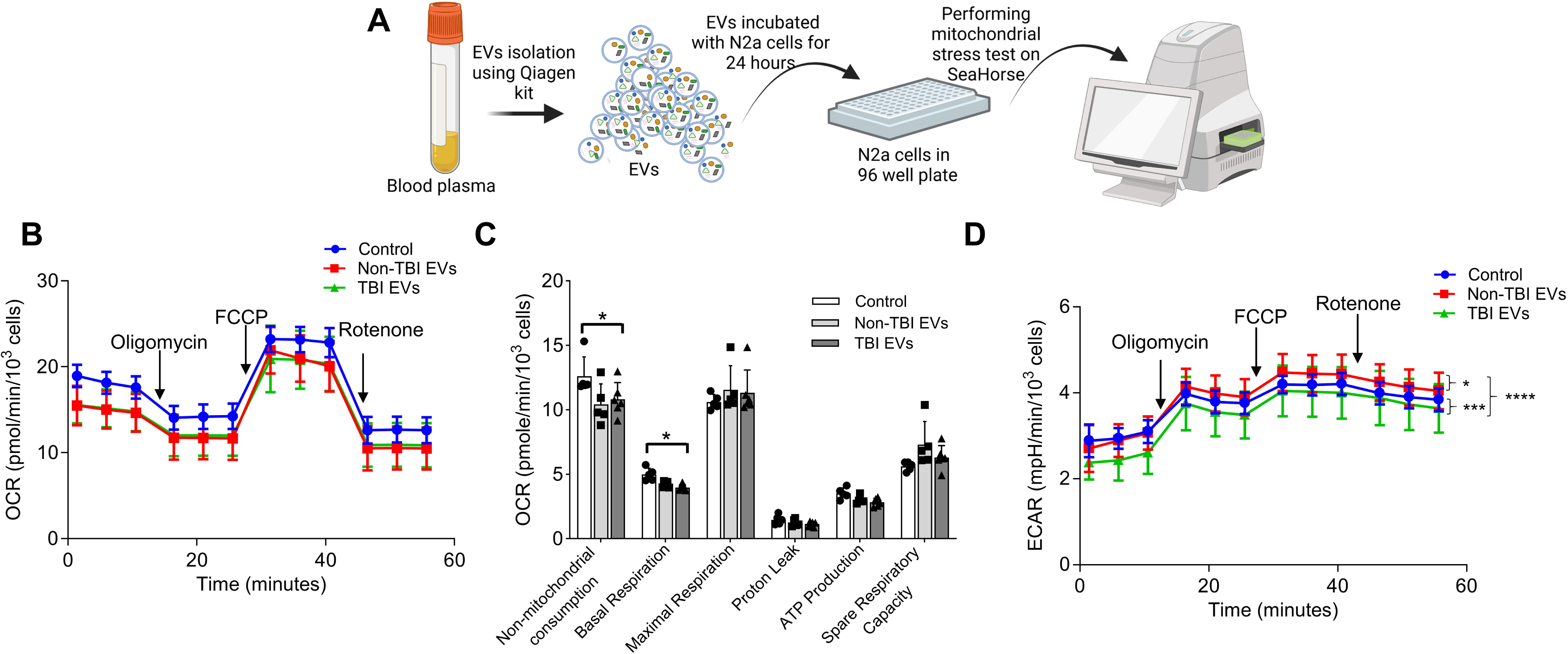
EVs from human TBI plasma affect glycolysis in N2a cells. (**A**) Schematic of EV isolation from human plasma, incubation of N2a cells without EVs (Control) or with TBI and non-TBI EVs, and Seahorse assay design. (**B–D**) Bioenergetic profiling of N2a cells incubated without EVs (control) or with human plasma-derived EVs from non-TBI or TBI subjects, analyzed using the Seahorse Flux Analyzer (Cell Mito Stress Test). Oxygen consumption rate (OCR) (**B** and **C**) are shown as lines (**B**) and box-and-whisker plot (**C**), and extracellular acidification rate (ECAR) (**D**) is shown as line. Mean ± SD, One-way ANOVA multiple comparison Tukey test was performed, *n* = 5 for control and non-TBI EVs and *n*=6 for TBI EVs, * *p* < 0.05, ** *p* < 0.01, *** *p* < 0.001, and **** for *p* < 0.0001.

## Discussion

One of the most exciting outcomes of plasma EV analysis is the discovery that a proportion of these vesicles originates in the brain.[14] The ability of EVs to cross the blood brain barrier (BBB) makes them suitable as markers for brain disease or injury. Beyond their diagnostic potential, the molecular cargo of brain-derived EVs may also provide critical insight into their functional roles in the underlying pathological processes.

Previous studies demonstrated that a subset of plasma EVs in patients with mild TBI originate from astrocytes and carry complement factors, pointing to an underlying neuroinflammatory process.[29] However, these studies did not analyze lipid content nor experimentally address the functional role of EVs in TBI pathogenesis. Building on our prior work implicating the sphingolipid ceramide and ceramide-binding proteins in the pathogenic activity of EVs in Alzheimer’s disease,[18, 24] we performed proteomic and targeted lipidomic analyses of plasma EVs from TBI patients as well as EVs from mice subjected to closed head injury (CHI), a complementary preclinical model for mild human TBI.[13, 28] Although differences between species and the predominance of severe TBI cases in the human cohort limit direct mechanistic interpretation, this combined approach offers a valuable translational link between clinical and experimental evidence for EV-mediated neuroinflammation. To directly assess EV functionality, we exposed neuronally differentiated N2a cells to TBI EVs and evaluated their impact on gene expression (transcriptomics) and mitochondrial function (Seahorse metabolic analysis). Our findings indicate that brain-derived EVs in plasma are not only biomarkers of neuroinflammation but also active contributors to the pathogenic processes triggered by TBI.

Previously, studies demonstrated the presence of GFAP in human plasma- and blood serum-derived EVs, supporting its potential as a TBI biomarker.[4, 52] Similarly, a sandwich ELISA identified GFAP and its breakdown products (GFAP-BDP) in TBI patient plasma.[57] Consistent with these reports, our findings showed a 25 kDa GFAP-BDP band by immunoblotting in both TBI plasma and plasma-derived EVs (Fig. 1D), correlating with disease severity (Supplementary Fig. 2). This represents a novel observation, highlighting GFAP-BDP as a sensitive circulating indicator of neuronal injury.

The proteomic screen focused on a protein band around 25 kDa clearly showing intense Ponceau (Fig. 1D) and Coomassie staining (Fig. 2A) when comparing TBI and non-TBI EVs from human plasma. We could not identify GFAP fragments, probably due to low abundance. However, immunoblot analysis confirmed the presence of 14-3-3 and CRP within plasma-derived TBI EVs (Fig. 2B and C). 14-3-3 proteins are highly expressed in the brain, particularly at synaptic membranes,[49] [32] While several 14-3-3 isoforms have been detected in plasma-derived EVs under non-neurological conditions, we demonstrate for the first time that 14-3-3 protein is elevated in plasma EVs from TBI patients, indicating a brain-derived contribution to the circulating EV pool, most likely due to 14-3-3 released within EVs due to neuronal injury. However, the specific contribution of 14-3-3 to TBI pathology remains to be investigated. Furthermore, although the isolation of brain-derived EVs from systemic EV populations remains technically challenging, CRP was detected exclusively in EVs isolated from TBI plasma, with no detectable signal in the EV-depleted fraction, indicating that its extracellular release is primarily EV-associated. The concurrent presence of a neuronal injury marker (GFAP) and an inflammatory mediator (CRP) exclusively within TBI blood plasma-derived EVs highlights a mechanistic link between central injury and inflammatory responses, raising the possibility that EV-associated GFAP and CRP could serve as clinically informative biomarkers capable of refining severity assessment in TBI.

Neuronal injury and neuroinflammation are well-established effects of TBI[15, 44, 63, 78] and are both closely linked to ceramide,[5, 19, 22] a bioactive lipid implicated in EV biogenesis and function.[61, 70] While increased ceramide concentrations have been observed in both the brain[5] and blood plasma[50] following TBI, the specific alterations in ceramides associated with blood-derived EVs have not yet been characterized. Our findings indicate that EV-associated ceramide may serve as a novel and minimally invasive biomarker with potential clinical relevance in TBI. Based on this, we assessed ceramide levels in EVs isolated from the plasma of TBI patients and controls. In human plasma, we observed increased levels of long-chain ceramides, particularly C18:1 and C20:0, with preferential association to CD9-positive EVs (Fig. 3). In our preclinical TBI mouse model, we observed a significant accumulation of ceramides, including C16:0, C18:0, C20:1, and C22:1 species, in the brain cortex (Fig. 4A). Furthermore, analysis of EVs isolated from TBI mouse brains revealed significantly increased levels of ceramide species C16:0, C18:0, C20:0, and C22:0, along with elevated levels of sphingolipid precursors, including dihydrosphingosine and sphingosine (Fig. 4B).

Although the mechanisms underlying elevated ceramide levels in TBI remain unclear, ASM, a key enzyme in ceramide generation, plays an important role in TBI pathology.[54, 55] We observed upregulation of ASM protein in mouse cortex with increased ceramide in TBI EVs (Fig. 5A and Fig. 4B), though ASM levels in plasma EVs did not differ between TBI and controls (Fig. 5C). These results suggest ASM-dependent ceramide accumulation within brain tissue, with ceramide subsequently incorporated into circulating EVs. Notably, TBI triggered ASM translocation to ependymal cilia (Fig. 5B), indicating localized ceramide generation and vesicle shedding. Consistent with TBI-induced ciliary loss due to vesicle shedding, human plasma EVs showed elevated levels of the ciliary marker Arl13b (Fig. 5D) and shortened ependymal cilia (Fig. 5B).[75] These findings suggest that a subset of ceramide-rich EVs originates from cilia, linking ASM redistribution to altered EV biogenesis. The cilia-to-EV pathway represents an underexplored communication axis with broad implications for TBI pathogenesis.

Mitochondrial dysfunction is a key feature of TBI.[31, 43, 62, 71] Transcriptomic analysis of N2a cells exposed to brain-derived TBI EVs indicated altered gene expression regulating oxidative phosphorylation, mitochondrial translation and sphingolipid metabolism, suggesting metabolic reprogramming (Fig. 6C). VDAC, a ceramide-binding mitochondrial apoptosis mediator,[20, 46, 77] was elevated in TBI cortex, consistent with our prior findings linking ceramide-rich EVs to mitochondrial dysfunction and neuronal cell death in Alzheimer’s disease.[24]

Our findings indicate that TBI induces a distinctive form of metabolic dysregulation in neurons that is not captured by changes in mitochondrial respiration (OCR), which remains largely unaltered (Fig. 7B). Instead, the disruption occurs at the interface of glycolysis and mitochondrial regulation. Figure 8 illustrates a working model in which ceramide-rich EVs generated after TBI impair neuronal glycolysis. Under physiological conditions, hexokinase 1 (HK1) is the predominant isoform in neurons,[73] linking glycolysis to mitochondrial ATP supply through binding to VDAC1/2 at the outer mitochondrial membrane.[2, 7] Following TBI, HK2 expression is induced in the injured cortex and hippocampus, indicating HK2 upregulation in the CNS after injury.[79] Previous studies suggest that ceramide can integrate into the outer mitochondrial membrane,[21, 64, 65] where it alters VDAC1/2 conductance and disrupts HK2–VDAC1/2 interaction. Displacement of HK2 from VDAC1/2 has been shown to induce cell death, uncouple glycolysis from mitochondrial ATP production, and reduce glycolytic capacity (lower ECAR) without overt suppression of oxidative phosphorylation.[23, 45, 47, 58]

**Figure 8.**
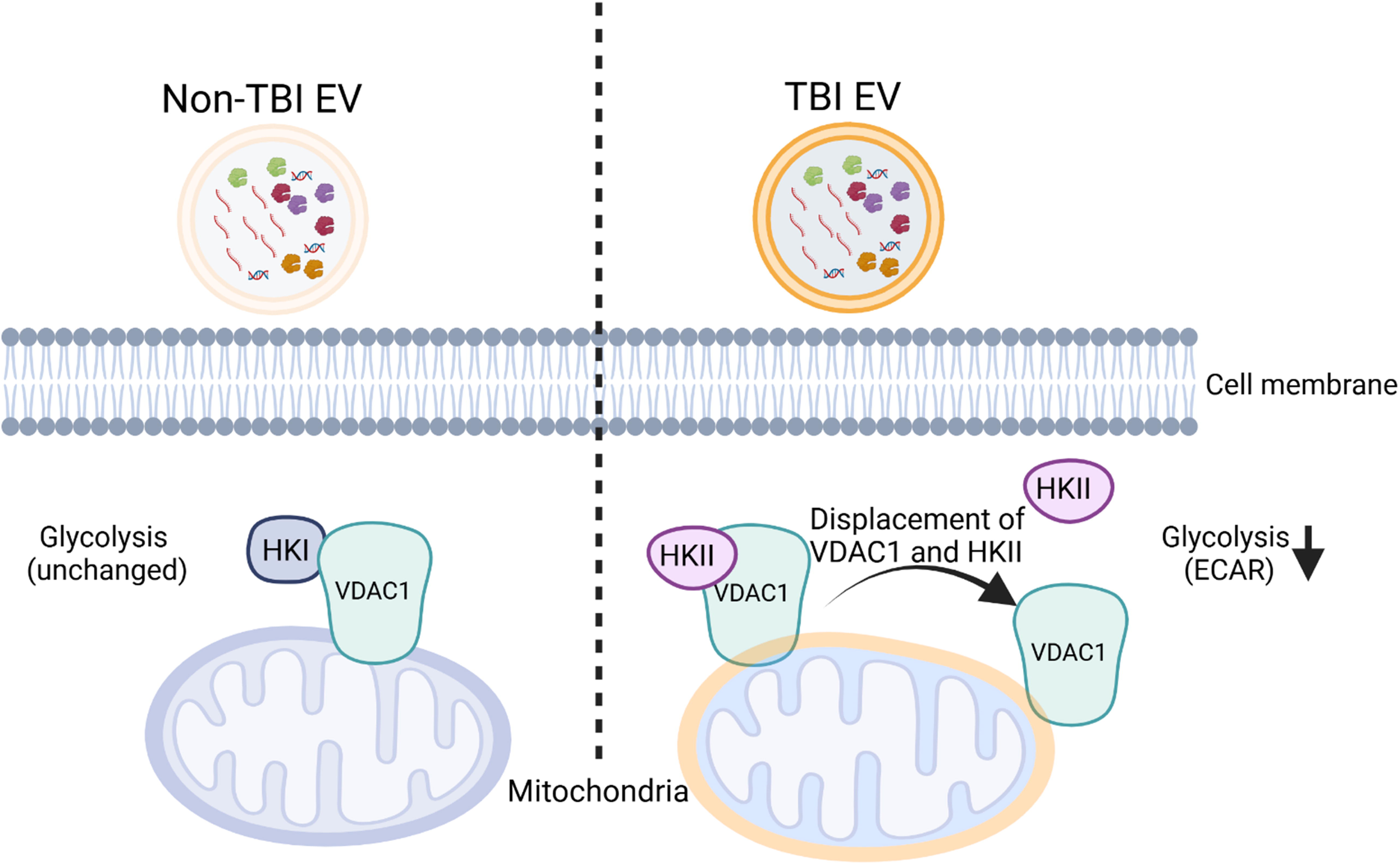
Proposed mechanism for the effect of ceramide-rich EVs on mitochondria and glycolysis. In non-TBI conditions (left), neurons predominantly express HKI, and exposure to control EVs has no significant impact on glycolytic activity. In contrast, following TBI (right), neurons predominantly express HKII, and exposure to ceramide-rich TBI EVs disrupts the HKII–VDAC1 interaction, leading to reduced glycolysis.

Consistent with this model, Seahorse analyses showed no significant change in overall OCR. However, ECAR, a readout of glycolytic flux, was significantly reduced in neurons exposed to human TBI EVs, indicating disrupted glycolytic metabolism. Compared with sham EVs, TBI EVs from mouse brain induced a modest but significant increase in cell death as measured by LDH release. In parallel, we observed elevated C16:0-ceramide in mouse brain lysates and brain-derived EVs (Fig. 4), together with increased p53 protein, a C16:0-ceramide–binding factor that promotes mitochondrial apoptosis.[26] Elevated p53 aligns with prior reports of p53 gene induction in injured brain regions after TBI.[51] Collectively, our findings suggest a mechanism whereby ceramide-rich EVs mediate neuroinflammatory and metabolic signaling through distinct cargo components, which may serve as potential biomarkers and contribute to neurotoxicity driving secondary injury. While the precise mechanisms by which TBI EVs may target neuronal mitochondria remain to be elucidated, ceramide-enriched EVs are proposed to disrupt HK–VDAC interactions, impair glycolytic coupling, and increase neuronal susceptibility to mitochondrial apoptosis, thereby amplifying metabolic dysfunction and secondary injury after TBI. Modulation of ceramide metabolism, inhibition of ASM activity, or blockade of EV release may mitigate these downstream effects, positioning ceramide-rich EVs as both mechanistic biomarkers and potential therapeutic targets in TBI.

## Conclusion

We show that human and experimental TBI generate ceramide-rich and inflammation-associated EVs that induce mitochondrial stress and neurotoxicity in neurons. Their molecular composition reflects both inflammatory activation and brain-derived origin. These data position ceramide-rich EVs as mechanistic contributors to TBI pathology and as potential targets for biomarker and therapeutic development.

## List of Abbreviations

ASM: acid sphingomyelinase
ApoA1: Apolipoprotein A1
BBB: blood–brain barrier
CNS: central nervous system
CRP: C-reactive protein
CHI: closed head injury
DEGs: differentially expressed genes
ECAR: extracellular acidification rate
EVs: extracellular vesicles
GFAP: Glial Fibrillary Acidic Protein
GCS: Glasgow Coma Scale
HK1: hexokinase 1
FcR: human Fc receptor
HSA: Human Serum Albumin
IP: immunoprecipitation
IACUC: Institutional Animal Care and Use Committee
LDH: lactate dehydrogenase
LC-MS/MS: liquid chromatography–tandem mass spectrometry
NTA: nanoparticle tracking analysis
OCR: oxygen consumption rate
SEC: size exclusion chromatography
Sph-1P: sphingosine-1-phosphate
TBS-T: Tris-buffered saline
VDAC1: voltage-dependent anion channel 1

## Declarations

### Ethics statement

Animal experiments were conducted in accordance with institutional guidelines and approved by the University of Kentucky Institutional Animal Care and Use Committee (IACUC; AUP #2017-2677).

### Consent for publication

Not applicable

### Availability of data and materials

The data that support the findings of this study are available from the corresponding authors, upon reasonable request.

### Author’s contributions

Erhard Bieberich and Zainuddin Quadri conceived the study and designed the experiments, including EVs isolation and characterization, cell culture assays, and fluorescence microscopy. Zainuddin Quadri, Zhihui Zhu, and Xiaojia Ren contributed to transcriptomics data interpretation and analysis. Zainuddin Quadri, Liping Zhang, and Simone Crivelli performed animal surgeries and collected brain and blood samples for downstream analyses. P.D. Kunjadia and Patrick Sullivan conducted the Seahorse assays in their laboratory. Bibi Broome and Tritia R. Yamasaki provided human plasma samples and associated clinical information. Zainuddin Quadri and Erhard Bieberich wrote the manuscript. All authors reviewed and critically revised the manuscript and approved the final version for submission.

## Acknowledgements

We thank the University of Kentucky NeuroBank, under the Neuroscience Research Priority Area, for providing human samples, and the Department of Physiology at the University of Kentucky College of Medicine (Chair: Dr. Alan Daugherty) for institutional support. The authors gratefully acknowledge the resources and support provided by the Proteomics and Mass Spectrometry Core Facility at Augusta University (Augusta, GA under supervision of Dr. Wenbo Zhi) and the Lipidomics Shared Resources Facility at the Medical University of South Carolina.

