## Supplemental Data for "Ceramide-rich extracellular vesicles as pathogenic biomarkers in traumatic brain injury"

**Supplemental figure and legends**

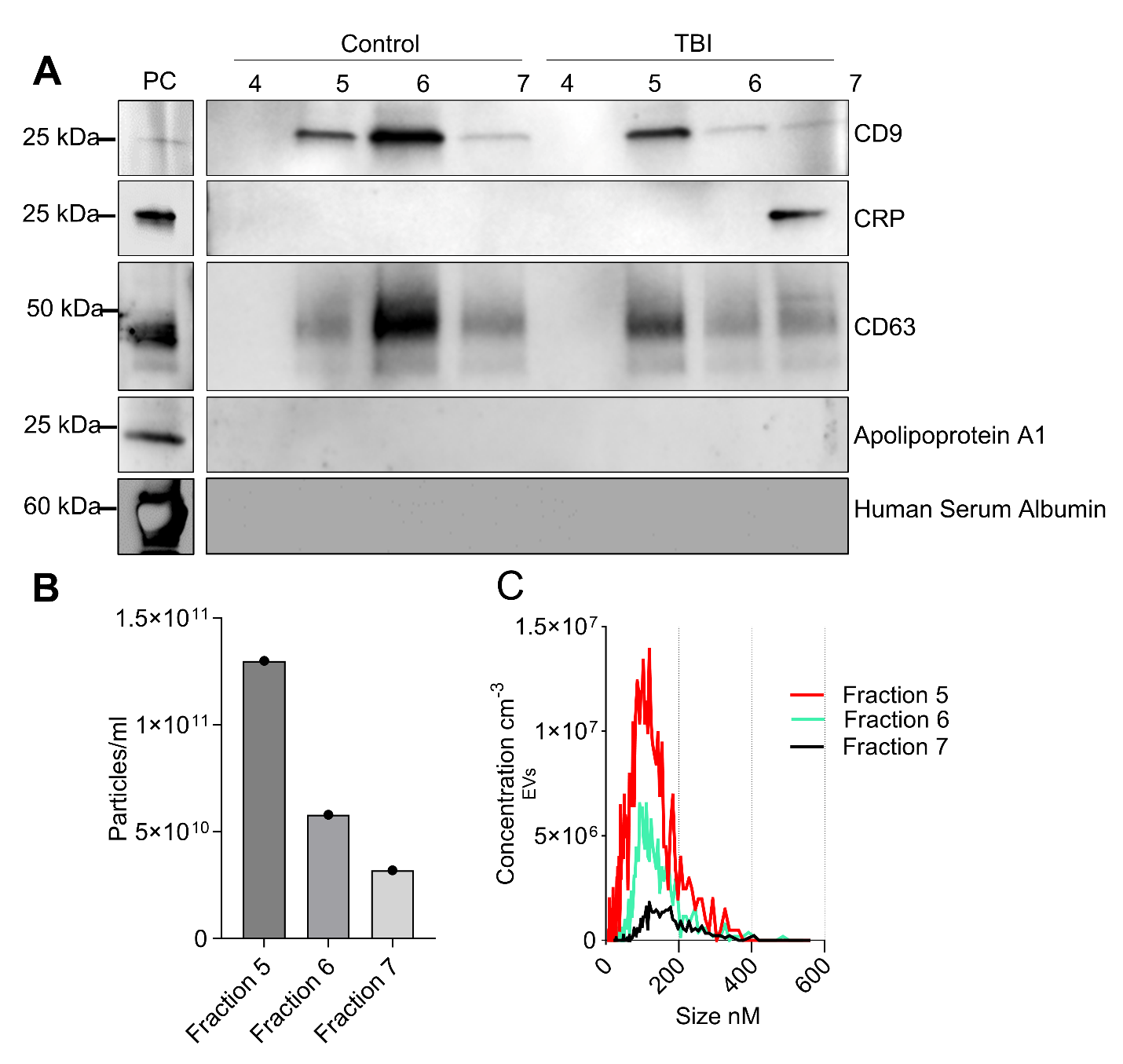
**Supplementary Figure 1. Isolation and characterization of plasma-derived EVs from TBI and control samples using SEC (iZon column).** (**A**) Immunoblot of EV-associated proteins CD9 and CD63, inflammatory protein CRP, and negative EV markers ApoA1 and human serum albumin. Human plasma was used as positive control (PC). (**B**) EV concentration (particles/ml) in fractions 5–7. (**C**) Particle size distribution curves of EVs isolated from fractions 5–7.

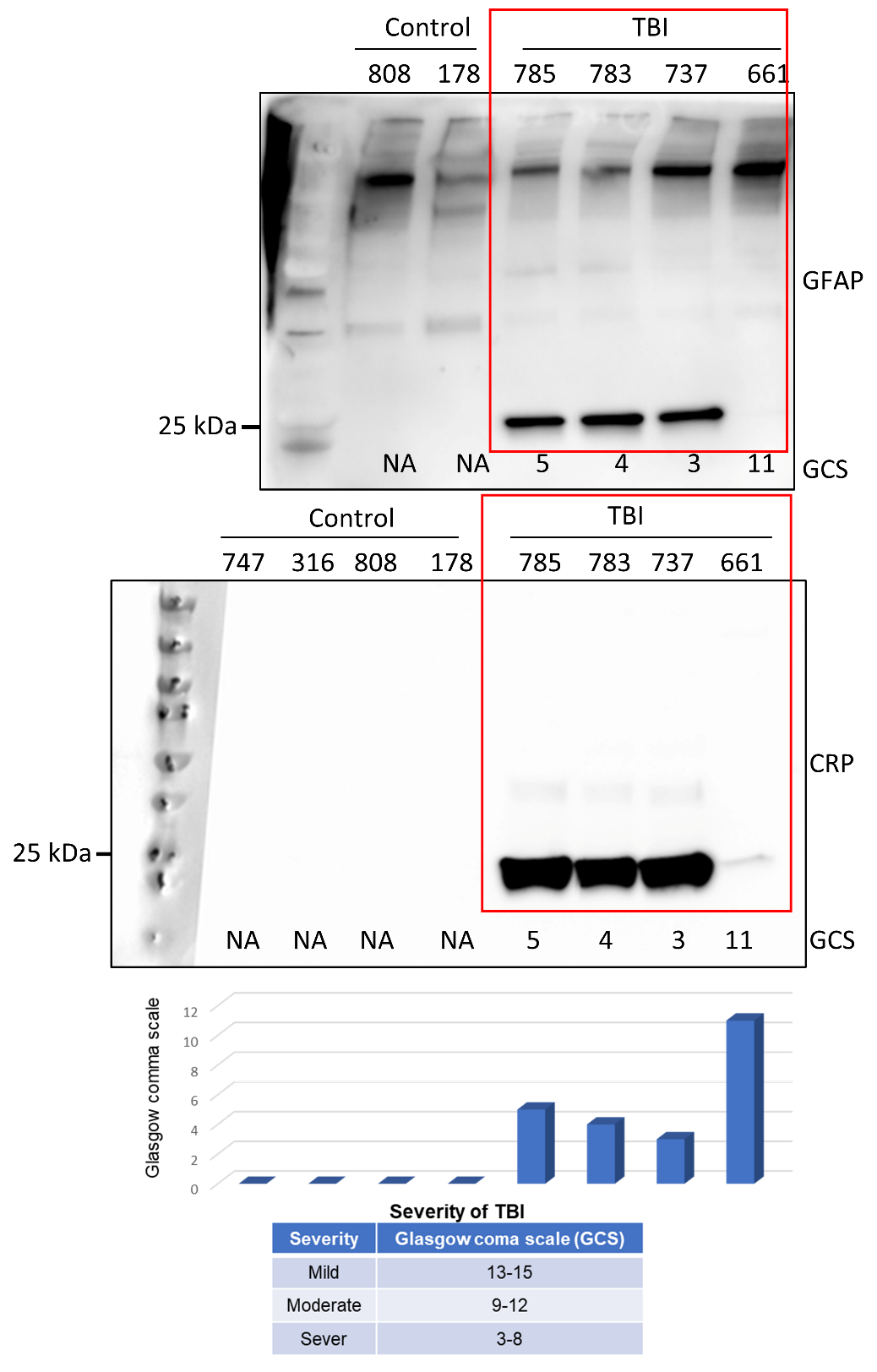

**Supplementary Figure 2. GFAP fragment and CRP levels correlate with TBI severity and inversely with Glasgow Coma Scale (GCS) score.** Immunoblot of GFAP-BDP (top) and CRP (bottom) in TBI EVs. Signals inversely correlate with GCS scores. Samples are numbered on top of the blots indicating patient ID; red frames indicate identical samples across blots. The lower panel presents a graphical representation of GCS scores corresponding to each TBI sample and NA (not applicable) denotes GCS for non-TBI.

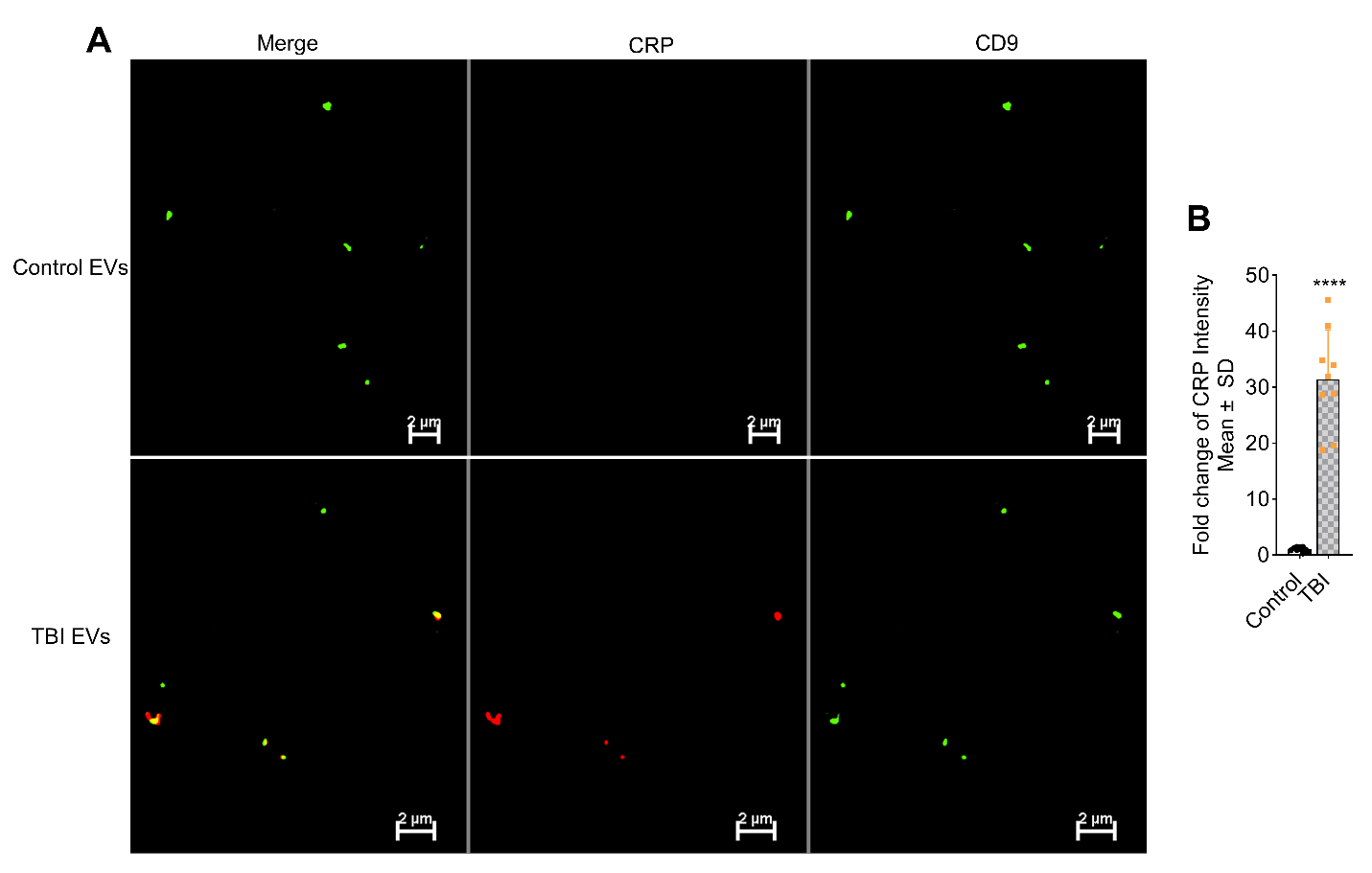
**Supplementary Figure 3. Immunofluorescence detection of EV-associated proteins.** (**A**) Human plasma-derived EVs immobilized on coverslips and immunolabeled for CRP and CD9. (**B**) Quantification showing fold-change of CRP signal in TBI EVs relative to controls. Mean ± SD, unpaired t-test, *n* = 10 for control and *n*=9 for TBI, * *p* < 0.05, ** *p* < 0.01, *** *p* < 0.001, and **** for *p* < 0.0001.

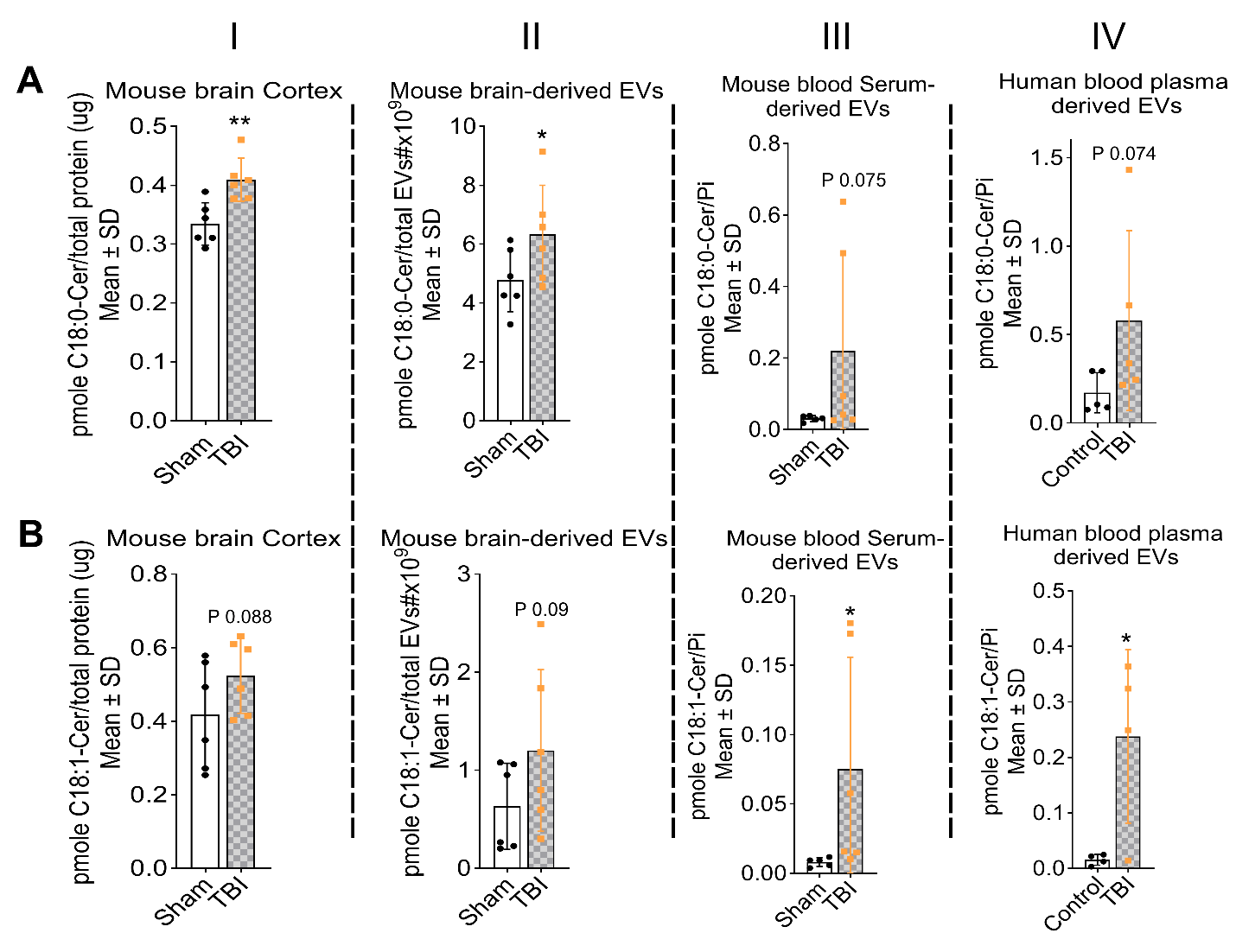
**Supplementary Figure 4. Levels of C18:0 and C18:1 ceramide species in TBI EVs.** Concentrations of C18:0 (**A**) and C18:1 (**B**) in mouse brain cortex (I), mouse brain-derived EVs (II), mouse serum-derived EVs (III) from TBI and sham groups, *n* = 6 for sham and TBI, as well as in plasma-derived EVs from human TBI and control samples (IV), *n* = 5 for control and *n*=6 for TBI. Mean ± SD, unpaired t-test, * *p* < 0.05, ** *p* < 0.01, *** *p* < 0.001, and **** for *p* < 0.0001. The values are normalized either to the total protein, number of EVs or lipid phosphate (Pi) as indicated.

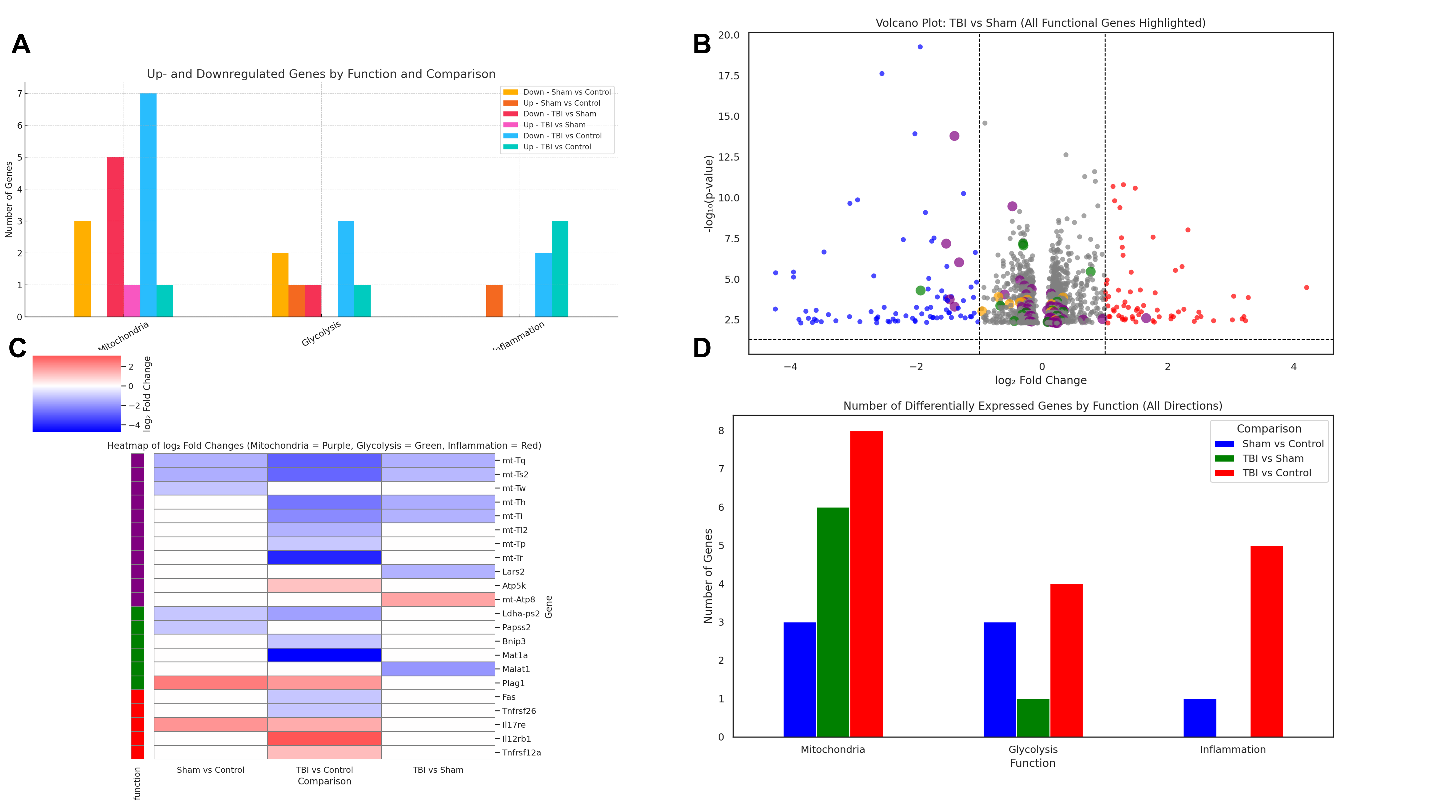
**Supplementary Figure 5. Transcriptomic changes in N2a cells after incubation with brain-derived EVs from TBI and sham mice.** (**A**) Bar graph showing significant number of up- and downregulated genes associated with mitochondrial, glycolytic, and inflammatory pathways across experimental groups. (**B**) Volcano plot highlighting mitochondrial (purple), glycolytic (green), and inflammatory (orange) genes. (**C**) Heat map of significant expressed genes associated with mitochondria (purple, top), glycolysis (green, middle) and inflammation (red, bottom) across experimental groups as indicated. (**D**) Number of significantly expressed genes in each pathway. Experimental comparisons include sham vs. control, TBI vs. sham, and TBI vs. control.

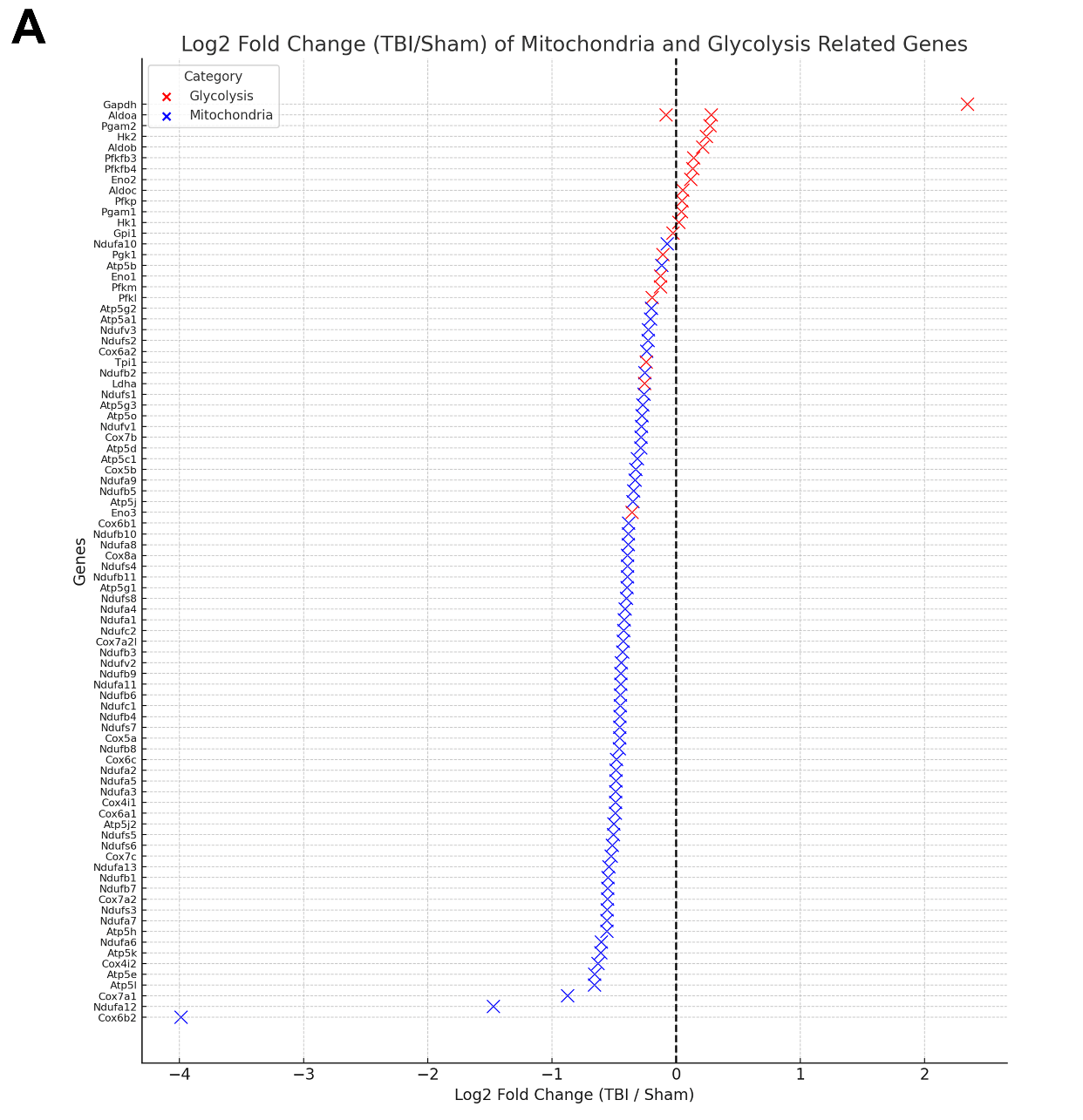
**Supplementary Figure 6. Transcriptomic analysis of glycolysis and mitochondrial-related genes in N2a cells incubated with EVs derived from mouse TBI brain.** (**A**) A volcano style scatter plot to show log₂ fold change on x-axis and lists genes related to mitochondria (blue) and glycolysis (red) on y-axis.

**Supplementary Table 1. List of proteins of approximately 25 kDa detected in extracellular vesicles (EVs) from TBI patients and control subjects.** Proteins were identified using mass spectrometry and are classified as unique to TBI EVs, unique to control EVs, or shared between the two groups. Each protein is annotated with its corresponding accession number.

| **Shared proteins (control and TBI)** | | | **Unique to control** | | | **Unique to TBI** | | |
| --- | --- | --- | --- | --- | --- | --- | --- | --- |
| **Accession No.** | **Description** | **MW [kDa]** | **Accession No.** | **Description** | **MW [kDa]** | **Accession No.** | **Description** | **MW [kDa]** |
| P02647 | Apolipoprotein A-I | 30.8 | P61026 | Ras-related protein Rab-10 | 22.5 | P02741 | C-reactive protein | 25 |
| A0M8Q6 | Immunoglobulin lambda constant 7 | 11.2 | A0A2R8Y619 | Histone H2B type 2-E1 | 13.5 | P0DOX7 | Immunoglobulin kappa light chain | 23.4 |
| P0CG04 | Immunoglobulin lambda constant 1 | 11.3 | P62249 | 40S ribosomal protein S16 | 16.4 | P63104 | 14-3-3 protein zeta/delta | 27.7 |
| P01834 | Immunoglobulin kappa constant | 11.8 | Q96JY6 | PDZ and LIM domain protein 2 | 37.4 | P02747 | Complement C1q subcomponent subunit C | 25.8 |
| P02649 | Apolipoprotein E | 36.1 |  |  |  | O00194 | Ras-related protein Rab-27B | 24.6 |
| P02743 | Serum amyloid P-component | 25.4 |  |  |  | O95755 | Ras-related protein Rab-36 | 36.3 |
| Q5VSP4 | Putative lipocalin 1-like protein 1 | 17.9 |  |  |  | O60258 | Fibroblast growth factor 17 | 24.9 |
| P69905 | Hemoglobin subunit alpha | 15.2 |  |  |  |  |  |  |
| P68871 | Hemoglobin subunit beta | 16 |  |  |  |  |  |  |
| A0A0C4DH25 | Immunoglobulin kappa variable 3D-20 | 12.5 |  |  |  |  |  |  |
